# Neocortical layer 4 in adult mouse differs in major cell types and circuit organization between primary sensory areas

**DOI:** 10.1101/507293

**Authors:** F. Scala, D. Kobak, S. Shan, Y. Bernaerts, S. Laturnus, C.R. Cadwell, L. Hartmanis, E. Froudarakis, J. Castro, Z.H. Tan, S. Papadopoulos, S. Patel, R. Sandberg, P. Berens, X. Jiang, A.S. Tolias

## Abstract

Layer 4 (L4) of mammalian neocortex plays a crucial role in cortical information processing, yet a complete census of its cell types and connectivity remains elusive. Using whole-cell recordings with morphological recovery, we identified one major excitatory and seven inhibitory types of neurons in L4 of adult mouse visual cortex (V1). Nearly all excitatory neurons were pyramidal and all somatostatin-positive (SOM^+^) non-fast-spiking neurons were Martinotti cells. In contrast, in somatosensory cortex (S1), excitatory neurons were mostly stellate and SOM^+^ neurons were non-Martinotti. These morphologically distinct SOM^+^ interneurons corresponded to different transcriptomic cell types and were differentially integrated into the local circuit with only S1 neurons receiving local excitatory input. We propose that cell-type specific circuit motifs, such as the Martinotti/pyramidal and non-Martinotti/stellate pairs, are optionally used across the cortex as building blocks to assemble cortical circuits.

## Main

The mammalian sensory neocortex is organized as a six-layer structure. In this stereotypical architecture, layer 4 (L4) is the main target of sensory inputs coming from the thalamus, thus acting as the first level of cortical processing for sensory signals. Understanding how L4 implements its computations requires to characterize the diversity of its constituent neuronal components and the connectivity among them.

Most previous studies of L4 have focused on primary somatosensory cortex (S1) of young rats and mice. Spiny stellate cells have been reported to be the dominant excitatory cell type, both in rat ^1–5^ and in mouse ^6^ (as a result of sculpting of initially pyramidal neurons during development ^7,8^). In contrast, inhibitory interneurons are highly diverse in terms of their genetic markers, morphologies and electrophysiological properties ^9^. Previous studies have reported three types of fast-spiking (FS), parvalbumin-positive (PV^+^) interneurons ^10^ and five types of non-FS interneurons ^11^, all of which have distinct morphologies. Several recent studies revealed that the somatostatin-positive (SOM^+^) interneurons form a single morphological population that has been called non-Martinotti cells ^12^ since their axons mainly target L4 ^13,14^, in contrast to typical Martinotti cells, which target L1. Interneuron types exhibit type-specific connectivity patterns. For example, PV^+^ FS interneurons receive strong thalamic inputs ^15–19^ while SOM^+^ non-FS interneurons receive weaker inputs ^20,21^. Both groups are reciprocally connected to local excitatory neurons and between each other ^10,14,16,18,22^, but PV^+^ inhibit each other while SOM^+^ do not ^23^.

Since most of these detailed studies were performed in S1 of young animals, it is unclear whether the cellular components of L4 and their connectivity profile are the same in adult animals and in other cortical areas. Recent large-scale studies of transcriptomic cell types in mouse and human cortex showed that most interneuron types are shared between cortical areas while the excitatory types are predominantly area-specific ^24,25^. In line with this, it has been shown that excitatory cells in L4 of mouse and rat primary visual cortex (V1) are pyramidal ^26,27^, in contrast to L4 in S1. However, there has been no systematic comparisons of anatomical and electrophysiological properties as well as connectivity profiles between L4 of different cortical areas, leaving an open question about the similarity in their cellular components and circuitry.

To address this question, we compared the microcircuit organization of adult mouse V1 L4 with S1 L4. We performed a thorough census of the morphologically defined cell types in V1 L4 of adult mice (median age 71 days) using multi-cell simultaneous whole-cell recordings combined with *post-hoc* morphological recovery ^28^. We identified several key differences in the cellular composition of L4 in V1 compared to the previous literature on S1, which we verified using targeted recordings of certain cell types in S1 L4 of similarly-aged mice. In addition, we mapped some of the observed morphological cell types to a reference transcriptomic cell type atlas ^24^ using Patch-seq ^29–31^. We further investigated the local connectivity profiles in L4 of both V1 and S1, finding similarities as well as some important differences in their circuitry.

## Results

### Morphologically defined cell types in L4 of adult mouse visual cortex

We characterized the electrophysiological and morphological features of L4 neurons in V1 of adult mice (*n*=129, median age 71 days, interquartile range 62--85 days, full range 55--330 days, Fig. S1) using whole-cell patch-clamp recordings combined with morphological recovery (see Methods). Altogether, we recovered and analyzed the morphology of *n*=1174 neurons (578 excitatory, 596 inhibitory).

Out of the 578 excitatory cells, 573 (99.1%) were pyramidal neurons (PYR), characterized by apical dendrites extending into layer 1 (L1), consistent with previous reports in rats ^26^ and young mice ^27^. These neurons did not show a complex arborization in L1, differing from typical layer 5 (L5) pyramidal neurons which generally have a prominent tuft in L1^32–34^ (Fig. 1a). Only five (0.9%) of the excitatory neurons were classified as spiny stellate cells based on the absence of the apical dendrites extending out of L4 to supragranular layers. These stellate cells had symmetrical non-polarized dendritic structure ^5^. The prevalence of PYRs among excitatory neurons in L4 of V1 was further supported by the fact that all labeled neurons recorded in Scnn1a-Cre/Ai9 mice (*n*=5), in which excitatory neurons in L4 are selectively labeled ^35,36^, were morphologically confirmed as PYRs (100%, 30/30). In terms of electrophysiology, PYRs exhibited large action potential (AP) width, high AP amplitude, and shallow afterhyperpolarization (AHP) which clearly discriminate them from GABAergic interneurons (Fig. 1b).

**Figure 1:**
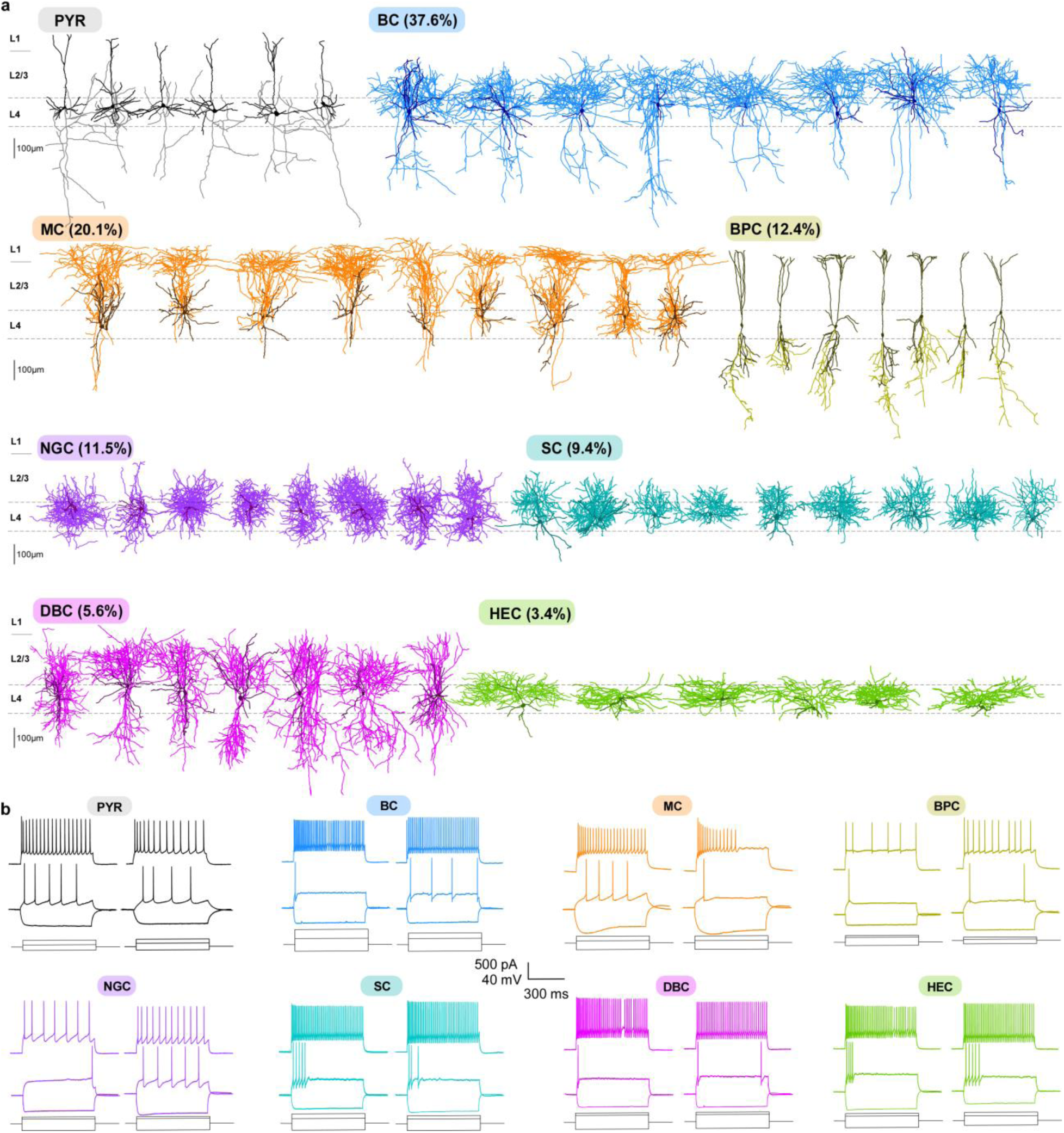
Morphological cell types in V1 L4. **(a)** Representative morphologies for each cell type. The dendrites are shown in a darker shade of color and the axons in a lighter shade. Types are sorted by abundance from high to low. Fractions indicate the proportion of inhibitory interneurons. PYR: pyramidal cells; BC: basket cells; MC: Martinotti cells, BPC: bipolar cells; NGC: neurogliaform cells, SC: shrub cells, DBC: double-bouquet cells, HEC: horizontally elongated cells. **(b)** Spiking responses to step currents for two exemplary cells from each of the eight morphologically defined cell types.

Interneurons showed a greater variability in both morphological and electrophysiological features. We used Viaat-Cre/Ai9 mice (*n*=47) to target GABAergic interneurons ^28,37^. Almost all labeled neurons recorded from these mice (95.5%, 234/245) were morphologically confirmed as interneurons, with only a small fraction (4.5%) of them being PYRs. On the other hand, all unlabeled neurons (*n*=133) were morphologically confirmed as excitatory neurons, suggesting that the entire population of interneurons in L4 was labeled in this Cre line. We identified seven GABAergic cells types (Fig. 1a) based on their morphology, following a widely used classification scheme based on the axonal geometry and projection patterns ^28,38–40^: basket cells (BCs; 37.6%, 88/234), Martinotti cells (MCs; 20.1%, 47/234), bipolar cells (BPCs; 12.4%, 29/234), neurogliaform cells (NGCs; 11.5%, 27/234), shrub cells (SCs; 9.4%, 22/234), double-bouquet cells (DBCs; 5.6%, 13/234), and horizontally elongated cells (HECs; 3.4%, 8/234). These morphological types varied greatly in abundance and electrophysiological properties (Fig. 1b).

The most abundant interneuron type was BCs, with large somata and thick axons originating from the apical side and projecting towards L2/3; they exhibited a fast-spiking (FS) firing pattern with narrow AP width and high maximal firing rate. They were followed by MCs, characterized by an ascending axon that projected to L1 and by their large membrane time constant. BPCs had a small soma, dendrites extending to L1 and L5, an axon projecting mostly downward to L5, and an irregular firing pattern. NGCs were characterized by a very thin axon that highly ramified around the soma; they were late-spiking and had large AP width. The remaining three types were all FS: SCs had a thick axon branching locally around their soma; DBCs had a thick axon projecting towards L5 and upwards to L2/3; HECs had a thick axon spreading horizontally within L4. A more detailed description of morphological and electrophysiological properties of all interneuron types can be found in the Supplementary Information.

We also performed experiments using several other Cre lines (parvalbumin-expressing, PV-Cre/Ai9, *n*=31; somatostatin-expressing, SOM-Cre/Ai9, *n*=14; and expressing vasointestinal peptide, VIP-Cre/Ai9, *n*=8) to relate genetic markers with morphological cell types (Fig. S2). The majority of morphologically recovered PV-Cre^+^ neurons were BCs (77.3%, 126/163 in PV-Cre/Ai9) and the rest were SCs (9.2%, 15/163), DBCs (12.3%, 20/163), and HECs (1.2%, 2/163). The majority of SOM-Cre^+^ neurons were typical MCs (91.8%, 56/61 in SOM-Cre/Ai9), while a small fraction exhibited an FS firing pattern and their morphological features matched those of BCs (8.2%, 5/61), in agreement with a previous report that due to potential off-target recombination,∼5--20% of neurons labeled in SOM-Cre line are FS ^28,41,42^ and PV^+^/SOM^−^ at the protein level ^28,41^. All VIP-Cre^+^ neurons in V1 L4 were BPCs (100%, 26/26 in VIP-Cre/Ai9). We did not encounter any labeled NGCs in any of these three Cre lines.

To support our expert classification, we fully reconstructed a subset of neurons from each inhibitory type (*n*=92 in total) and trained a regularized logistic regression classifier to discriminate between each pair of inhibitory cell types (see Methods). We used 2D density maps and a set of morphometric statistics (Fig. S3) as predictors ^43^. Across all 21 pairs, the average cross-validated classification accuracy was 0.92, with most pairs discriminated almost perfectly (Fig. 2a, left). However, SC/HEC and SC/NGC pairs showed only ∼0.65 classification accuracy. Visualisation of this dataset with t-SNE (Fig. 2a, right) indicated that SC/HEC and SC/NGC types were partially overlapping, as well as BC/DBC. Overall, this analysis suggests that while most morphological classes can be very well discriminated, some may be partially overlapping. An important caveat is that low classification accuracy can also be due to an insufficient sample size.

**Figure 2:**
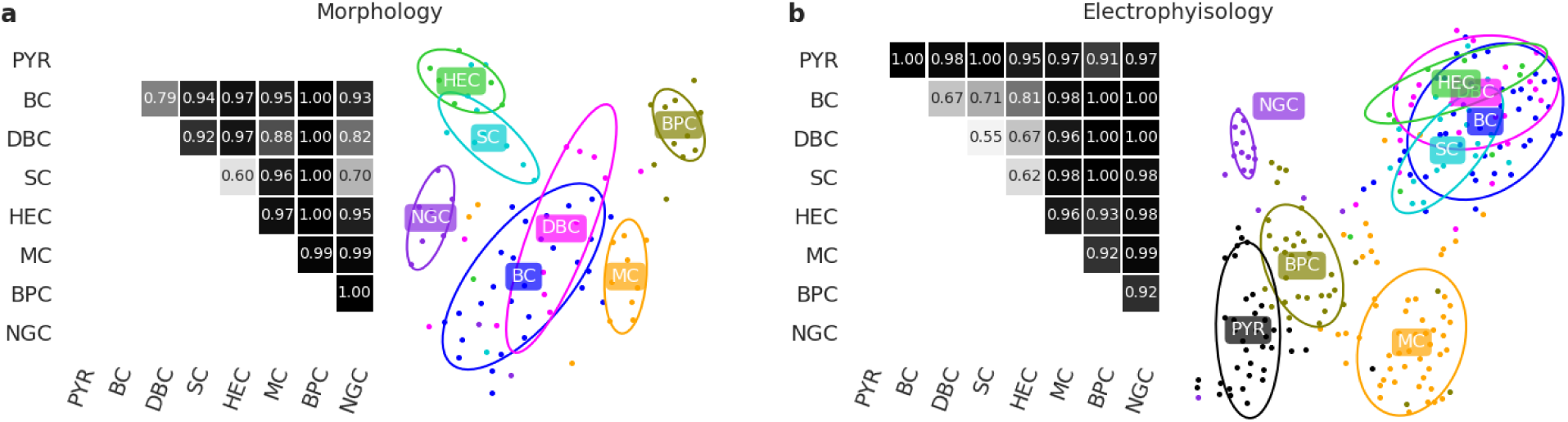
Discriminability of morphologically defined cell types in V1 L4 using morphological and electrophysiological properties. **(a)** Cross-validated pairwise classification accuracy for each pair of inhibitory cell types, using regularized logistic regression on a diverse set of morphological features. Total sample size *n*=92. Right: 2D visualisation of the same *n*=92 cells in the space of morphological features using t-SNE. Ellipses show 80% coverage assuming 2D Gaussian distributions and using robust estimates of the mean and the covariance (i.e. ellipses do not include outliers). **(b)** Cross-validated pairwise classification accuracy for each pair of cell types, using electrophysiological features. Total sample size *n*=235. Right: 2D visualisation of the same *n*=235 cells in the space of electrophysiological features using t-SNE.

To further explore variability in electrophysiological properties between cell types, we characterized the firing pattern of a subset of neurons (*n*=235) using 13 automatically extracted electrophysiological features (Fig. S4). Most features exhibited strong differences between cell types (Fig. S5). Two-dimensional visualisation of this dataset using t-SNE (Fig. 2b) showed that all four PV^+^ cell types overlapped in one group of electrophysiologically similar FS neurons, while the other four types (PYR, NGC, BPC, and MC) each had distinct firing patterns. We confirmed this using pairwise classification with regularized logistic regression (Fig. 2b): the average cross-validated classification accuracy between the FS types was only 0.67, while the average accuracy across all other pairs was 0.98.

### V1 differs from S1 in major L4 cell types

In contrast to V1 L4, stellate cells are known to be abundant in S1 L4 of rats and mice ^1–7^. To confirm this, we recovered L4 excitatory cells in S1 (*n*=24 mice, including *n*=5 Scnna1-Cre/Ai9) with the same method as in V1. We found that indeed 76.6% (85/111) of the recovered spiny neurons did not have a clear apical dendrite and were thus classified as stellate cells (Fig. 3B), while the remaining 23.4% were pyramidal cells. This confirms that, unlike in V1, stellate cells are the predominant excitatory population in L4 of adult mouse S1 (Fisher’s exact test for difference in the fraction of stellate cells between V1 and S1: p<0.0001; 85/111 vs. 5/578).

**Figure 3:**
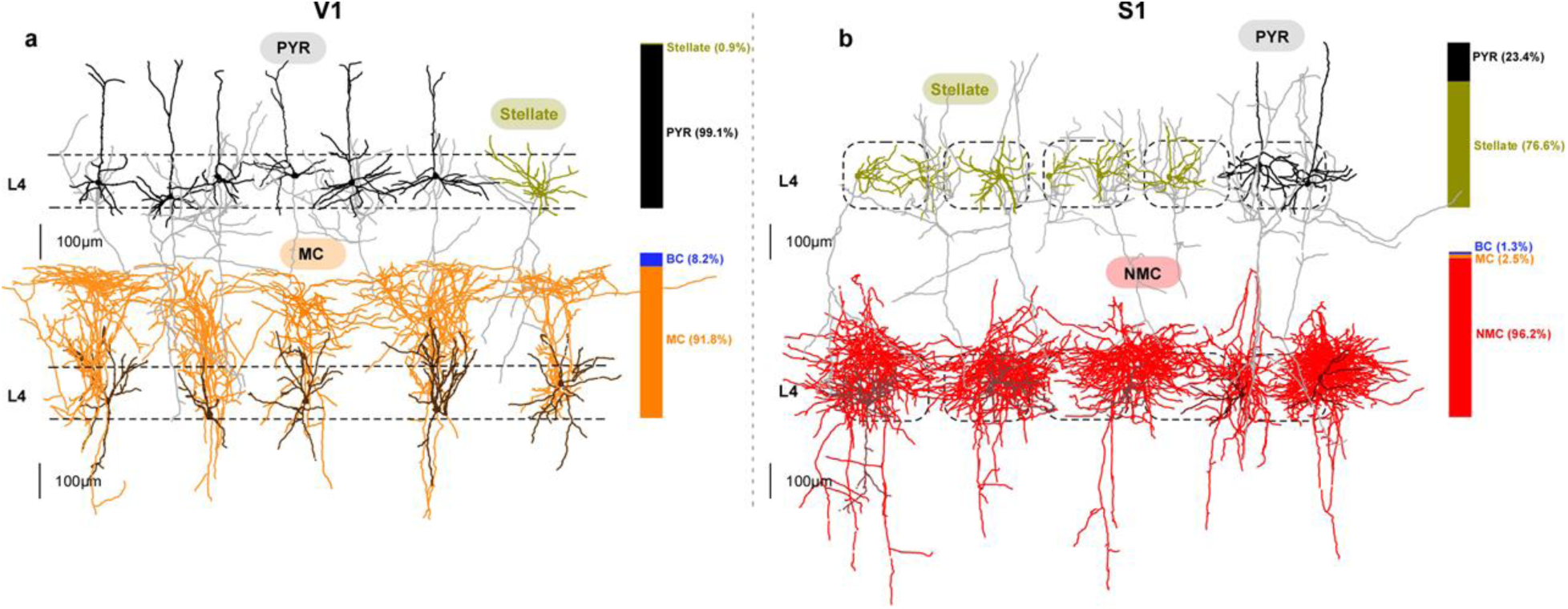
V1 differs from S1 in excitatory cells and SOM^+^ interneurons in L4. **(a)** Representative morphologies of excitatory and SOM^+^ neurons in V1 L4. Bar graphs indicate the fractions of each cell type among all morphologically recovered excitatory neurons (top) and all morphologically recovered SOM-Cre^+^ neurons (bottom). **(b)** The same in S1 L4. Dashed rectangles represent individual cortical barrels.

Recent evidence indicates that most, if not all, L4 SOM^+^ cells in mouse S1 are non-MC having axons mostly localized within L4, in stark contrast to typical MCs ^14^. Indeed, we found that in S1, almost all L4 SOM-Cre^+^ neurons we recovered (96.2%, 76/79, from *n*=19 SOM-Cre/Ai9 mice) had non-MC morphology characterized by a thin, highly ramifying axon mostly residing within L4 (Fig. 3B). Only two cells showed an ascending axon projecting to L1 typical of MCs (2.5%, 2/79) and one was characterized by a thick axon branching similarly to BC with a FS firing pattern (1.3%, 1/79) (Fisher’s exact test for difference in the fraction of NMCs between S1 and V1: p<0.0001; 76/79 vs. 0/61). We follow the convention of a previous study ^12^ and refer to these SOM^+^ neurons that dominate in L4 of S1 as non-Martinotti cells (NMCs).

The NMCs also differed in their firing pattern from MCs recorded in V1: they had a higher maximal firing rate, a lower AP width, and a lower membrane time constant (Fig. 4a and Fig. S5). This resembles the FS firing pattern, and one previous study called NMCs “quasi-FS” ^13^. Comparison of electrophysiological features between MCs, NMCs, and FS cells revealed that NMCs were “in between” MCs and FS cells in terms of their firing patterns and intrinsic membrane properties (Fig. S5, S6).

**Figure 4:**
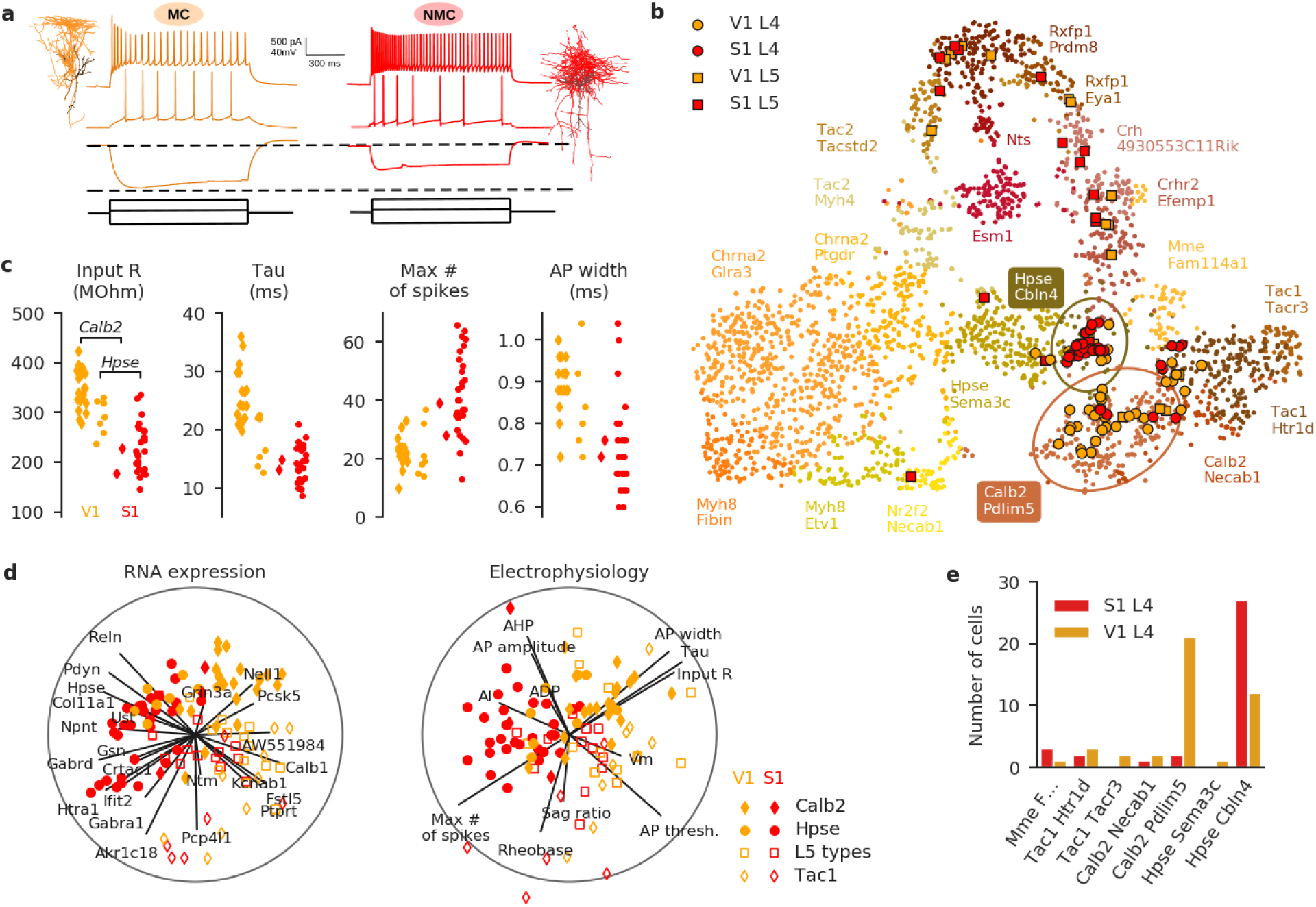
SOM^+^ interneurons in L4 of V1 and S1 differ in electrophysiological properties and transcriptomic profile, as shown by Patch-seq. **(a)** Morphologies and firing patterns of two exemplary cells, from V1 (orange) and S1 (red) respectively. **(b)** Mapping of the Patch-seq cells (*n*=110) to the t-SNE visualization of the transcriptomic diversity among *Sst* types from Tasic et al. ^24^ t-SNE was done on all cells from *Sst* types except for *Sst Chodl* that is very well separated from the rest (20 clusters; *n*=2701 cells), using 500 most variable genes (see Methods). Two ellipses show 90% coverage areas of the two types where the most Patch-seq cells land. Mapping to t-SNE was performed as we described elsewhere ^47^, see Methods. “Sst” was omitted from type names for brevity. **(c)** Four electrophysiological features that differed most strongly (Cohen’s *d*>1) between V1 L4 and S1 L4 cells. Only cells assigned to *Sst Calb2 Pdlim5* and *Sst Hpse Cbln4* types are shown. Note that the values are not directly comparable to those shown in Fig. S5 because Patch-seq experiments used a different internal solution compared to regular patch-clamp experiments without RNA extraction. **(d)** Sparse reduced-rank regression analysis ^46^: the left biplot shows two-dimensional projection in the transcriptomic space that is optimized to reconstruct the electrophysiological features. The right biplot shows the corresponding two-dimensional projection in the electrophysiological space; it should “match” to the left plot if the model is accurate. Color denotes brain area (orange for V1, red for S1), marker shape denotes transcriptomic type that each cell was assigned to (circles: *Hpse Cbln4* type; diamonds: *Calb2 Pdlim* type; open diamonds: three *Tac1/Mme* types and the neighbouring *Calb2 Necab1* type; open squares: all other types). Individual electrophysiological features and genes selected by the model are depicted with lines showing their correlations to the two components. Circles show maximal possible correlation. Cross-validated estimate of the overall R-squared was 0.16, and cross-validated estimates of the correlations between the horizontal and vertical components were 0.70 and 0.50 respectively. **(e)** Type assignments of the Patch-seq cells from L4.

To further investigate the differences between MCs in V1 and NMCs in S1, we used the Patch-seq ^29–31^ technique which combines patch-clamp recordings with single cell transcriptomics. Using *n*=6 SOM-Cre/Ai9 mice, we sequenced RNA of SOM-Cre^+^ neurons in L4 of V1 and S1 (*n*=42 in V1 and *n*=35 in S1 after quality control), and also in L5 of each area (*n*=17 and *n*=16 respectively). We obtained on average 1.1 million reads per cell (median; mean±SD on a log10 scale: 6.0±0.3) and detected 6.4±1.6 thousand (mean±SD) genes per cell (Fig. S7). We mapped these cells to a large transcriptomic cell type dataset ^24^ that contained 21 somatostatin types with 2880 neurons from V1 and ALM. The quality of the mapping was equally good for V1 and S1 cells (Fig. S7), suggesting that the V1+ALM dataset is a reasonable reference for S1 interneurons. This is in agreement with the idea that inhibitory transcriptomic cell types are shared across cortical regions ^24,25^. Three cells (excluded from the counts given above and from further analysis) had fast-spiking firing pattern, did not express SOM, and mapped to *Pvalb Reln Itm2a* transcriptomic type, likely corresponding to the basket cells that we found labeled in the SOM-Cre line (Fig. 3). All other cells mapped to *Sst* transcriptomic types.

Most L4 cells (81%, 62/77) were assigned to one of the two transcriptomic types: *Sst Calb2 Pdlim* and *Sst Hpse Cbln4* (Fig. 4B,E), with S1 cells falling almost exclusively into the *Hpse* type (27/29) and V1 cells falling preferentially into the *Calb2* type (21/33) (p<0.0001, Fisher’s exact test). This suggests that *Sst Calb2 Pdlim* is a MC type, in agreement with the conclusions of Tasic et al. ^24^ based on the data from Paul et al.^44^, and that *Sst Hpse Cbln4* is a NMC type, in agreement with Naka et al. ^45^. However, this raises the question of why some V1 L4 SOM^+^ cells, none of which had a NMC morphology (see above), had a NMC transcriptomic profile, both among our Patch-seq cells and in the Tasic et al. dataset ^24^.

To answer this question, we looked at electrophysiological features that were most different between SOM^+^ interneurons in V1 and S1 (Cohen’s *d*>1: input resistance, membrane time constant, maximum firing rate, and AP width) and found that for two of them (input resistance and membrane time constant) V1 cells belonging to the *Hpse* type had values more similar to the S1 cells than to the V1 cells from the *Calb2* type (Fig. 4d). This suggests that electrophysiologically, V1 *Hpse* MC cells are in between V1 *Calb2* MC cells and S1 NMC cells.

The relationship between gene expression and electrophysiological features can be visualized using the sparse reduced-rank regression analysis that we have recently introduced ^46^. This technique aims to reconstruct all the electrophysiological features using a two-dimensional projection of the expression levels of a small set of genes (Fig. 4d). The optimal number of genes was found using cross-validation (see Methods). This analysis supports our conclusion that V1 *Hpse* MCs are “in between” *Calb2* MCs and NMCs in terms of electrophysiology. Interestingly, this analysis also showed that some of the cells assigned to the *Tac1* and *Mme* types had a distinct fast-spiking-like firing pattern which was different from firing patterns of MCs and NMCs (but was not as sustained as the proper FS pattern). These three SOM^+^ transcriptomic cell types have recently been identified in Tasic et al. ^24^, and do not have known morphological or electrophysiological counterparts.

The L5 SOM^+^ cells that we sequenced in both areas mostly mapped to a different set of transcriptomic types than the L4 SOM^+^ cells, but there were no apparent differences between S1 and V1 in terms of transcriptomic cell types (Fig. 4b).

### Connectivity among excitatory and SOM^+^ neurons in L4 of V1 vs. S1

So far, we have described major differences in the morphology, electrophysiology, and transcriptomic signatures of excitatory neurons and SOM^+^ interneurons in L4 between V1 and S1. We next investigated whether there are differences in their connectivity profiles as well, using simultaneous multi-cell patch-clamp recordings. We found that certain connectivity patterns between them are very similar across both areas (Fig. 5). First, the connection probabilities among excitatory cells were low in both areas (1.0%, 7/701 in V1; 2.5%, 3/122 in S1). Second, the connection probabilities between SOM^+^ cells were also low in both areas (0%, 0/68 in V1; 3.8%, 2/52 in S1). Third, the connection probabilities from SOM^+^ cells to excitatory cells were high in both areas (21.1%, 30/142 in V1, 26.6%, 17/64 in S1). In addition, despite their low connectivity via chemical synapses, both MCs in V1 and NMCs in S1 were similarly often interconnected by gap junctions (MCs: 23.5%, 8/34; NMCs: 30.7%, 8/26; Fig. S6).

**Figure 5:**
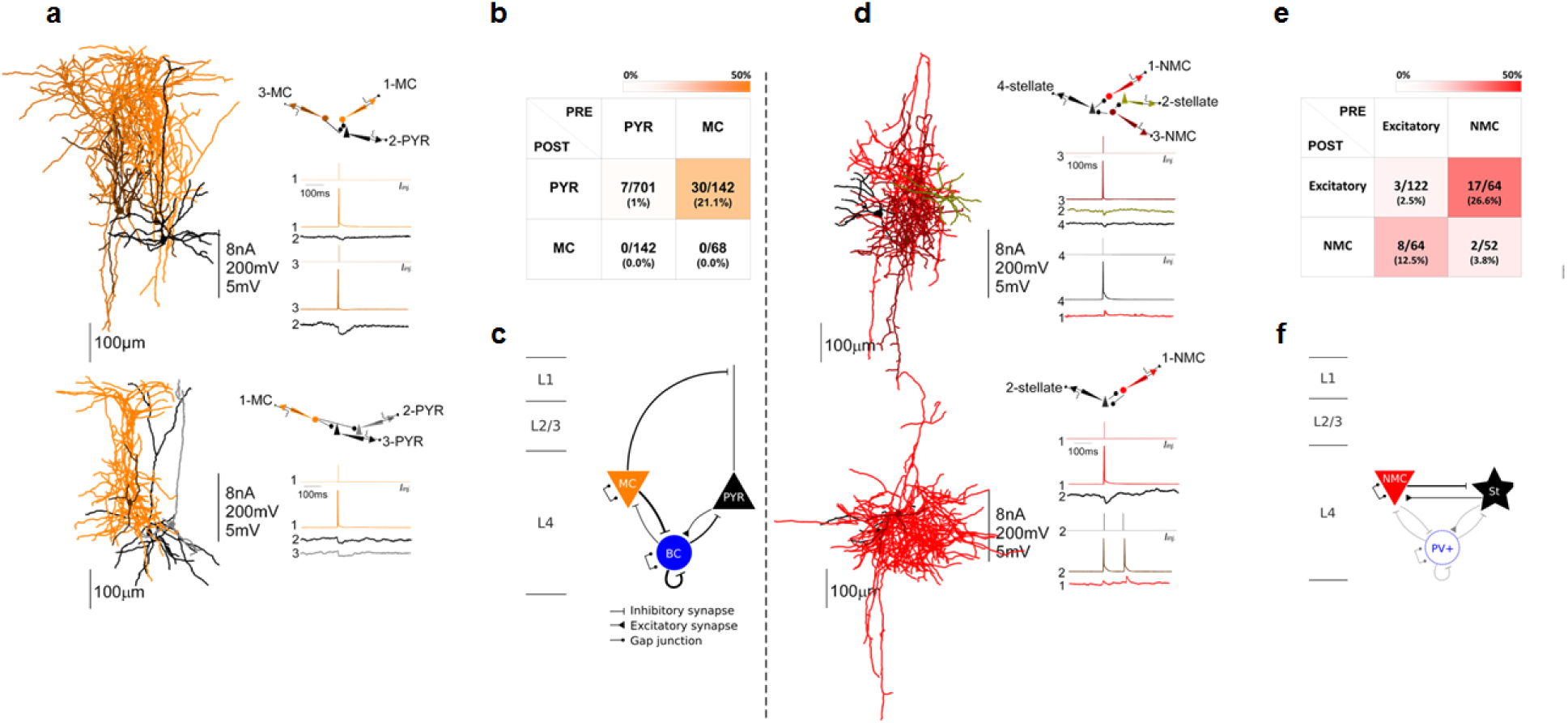
Connectivity between excitatory and SOM^+^ cells in L4 of V1 and S1. **(a)** Examples of simultaneous recordings from excitatory and SOM^+^ neurons in V1 L4. Recorded neurons were close to each other (generally less than 250μm). Vertical scale bar indicates: amplitudes of injected current in nA, amplitude of APs in mV and amplitude of uEPSP or uIPSP in mV. **(b)** Color-coded connectivity matrix shows the connection probability between cell types as a percentage of tested potential connections. Average of uEPSP and uIPSP as well as PPR are reported in Fig. S7. **(c)** Schematic of the local circuitry in L4 V1. For the connectivity involving BCs, see Fig. S8. For gap junctions, see Fig. S8. Line thickness corresponds to connection probability. **(d-f)** The same for L4 S1. In the schematic in panel (F), connectivity involving BCs is taken from Ma et al. ^23^ All these BCs connections are shown with the same strength as that study used juvenile (P15) mice and so connection strengths are not directly comparable to the values obtained in our experiments. Regarding gap junctions between FS interneurons see also ^49,55^.

On the other hand, we found a striking area-specific difference in connection probabilities from excitatory to SOM^+^ neurons. In S1, NMCs received facilitating synaptic connections from local excitatory neurons (12.5%, 8/64), in line with previous studies in young rodents ^14,16^. In contrast, we did not find any connections (0%, 0/142) from local excitatory neurons to MCs in V1 (*p*=0.0002, Fisher’s exact text). This was also in stark contrast to MCs in L2/3 and L5 of adult mouse V1, which receive strong facilitating synaptic inputs from local PYRs in the same layers ^28^ (see Discussion for further considerations).

In addition, we tested the connectivity in V1 L4 including BCs (Fig. S10). We found that BCs followed the same connectivity rules as previously found in other layers ^28,48^. PYRs connected to BCs with probability 12.5% (38/303), MCs inhibited BCs with probability 32.6% (15/46), and BCs inhibited each other (36.7%, 75/204), MCs (13.0%, 6/46) and PYRs (25.7%, 78/303). All of these connection patterns have also been reported in S1 L4 of young mice ^23^. We also found that BCs were electrically coupled to each other with probability 27.5% (28/102) but were never electrically coupled to MCs (0/23), in agreement with previous findings that gap junctions exist between inhibitory cells of the same type ^49^.

Notably, the connection probability between PYRs in V1 L4 was very low, consistent with our previous work in other layers in adult animals ^28^, but in contrast to the findings in young and juvenile rodents ^50,51^. To confirm that this low connectivity reflects an age effect, we measured the connectivity between PYRs in V1 L4 at different ages (P15-20 and P30-40, *n*=5 each) using Scnn1a-Cre/Ai9 mice. We found that the connection probability monotonically decreased with age (Fig. S11): from 13.2% in P15-20 (15/114) to 5.1% in P30-40 (8/156) to 1.0% (7/701) reported above for the P55+ mice with median age P71. This is in agreement with the recent study that found 6.3% (20/315) connection probability in V1 L4 of P47±6 mice ^52^.

When measuring connectivity in S1 L4, no special care was taken to ensure that the tested cells were within the same barrel. At the same time, it is known that cells in S1 L4 preferentially make intra-barrel connections ^2,3^. To address this concern, we performed a separate series of experiments using *n*=8 Scnn1a-Cre/Ai9 mice to test intra-barrel connectivity of excitatory neurons. We used the tdTomato fluorescence signal to detect the barrels during patch clamp recordings ^53^ and performed cytochrome oxidase staining in a subset of slices to confirm that the fluorescence signal reliably corresponded to barrel boundaries ^3^ (see Methods and Fig. S12). The measured connection probability was 5.2% (5/104) which was larger than the value reported above (2.5%, 3/122) but not significantly different from it (*p*=0.48, Fisher’s exact test). Both estimates are substantially lower than the existing estimates of intra-barrel connectivity obtained in young animals (30--35%) ^2,3,54^ which is in line with the decrease in local excitatory connectivity with age discussed above for V1 (see also Fig. S11).

## Discussion

### Morphological cell types in L4: V1 vs. S1

We described eight morphological cell types in L4 of primary visual cortex (V1) in adult mice as well as the connectivity patterns of three most abundant cell types. We found that nearly all excitatory neurons are pyramidal cells (as was previously described in rats ^26^, guinea pigs ^56^, and young mice ^27^), in stark contrast to L4 of S1 where the majority of excitatory neurons are stellate cells ^5,7,57^. Interestingly, L4 stellate cells in ferret V1 and mouse S1 develop postnatally from neurons that resemble pyramidal cells with an upward projecting apical dendrite ^7^. The near absence of stellate cells in V1 L4 of adult mice (as old as 11 months in our experiments) suggests a different developmental path in this case. Excitatory neurons in V1 L4 of other species such as cats ^58^ and monkeys ^59^ are also known to be stellate. It remains an open question, why pyramidal cells in rodent V1 L4 remain pyramidal, whereas L4 excitatory cells in rodent S1 and in V1 of other non-rodent species develop into stellate cells. We suggest one hypothesis below when discussing their circuit organization.

We found that all non-fast-spiking SOM^+^ neurons in V1 L4 are Martinotti cells (MCs), which is also in contrast to S1 L4 where almost all SOM^+^ neurons are non-Martinotti ^13,14^. Using Patch-seq, we showed that SOM^+^ MCs in V1 L4 and SOM^+^ NMCs in S1 L4 correspond to two different transcriptomic cell types (*Sst Calb2 Pdlim* and *Sst Hpse Cbln4* respectively) previously identified in a large-scale transcriptomic cell atlas ^24^.

We relied on manual expert classification to isolate the morphological types. Unlike in transcriptomics, where automatic unsupervised clustering is commonplace ^24^, morphological studies usually do not use it, because of low numbers of manually reconstructed neurons and multiple challenges to data analysis of morphological data. One recent study done in parallel to our work ^60^ attempted clustering of neural morphologies from all layers of adult mouse V1. There is a broad agreement between their types from L4 and our types. There are also some differences: they split abundant types (e.g. PYRs and BPCs) into multiple narrow sub-clusters, while at the same time missing some rare types such as HECs.

### Transcriptomic types of SOM^+^ interneurons in L4: V1 vs. S1

Although MCs and NMCs are morphologically distinct, with no ambiguous morphological forms, they form more of a continuum in both transcriptomic and electrophysiological space. In the Tasic et al. reference dataset ^24^, the MC and the NMC clusters (*Sst Calb2 Pdlim* and *Sst Hpse Cbln4* respectively), although distinct, were close and partially overlapping in the t-SNE visualisation (Fig. 4b). Consistent with this, Tasic et al. ^24^ also found intermediate cells between the “core” members of these two clusters. We showed that, electrophysiologically, MCs and NMCs also form a continuum (Fig. 4d, Fig. S6) with all electrophysiological features having unimodal distributions (Fig. 4c). This is in agreement with the findings of Naka et al. ^45^ who demonstrated an electrophysiological continuum between NMCs in S1 L4 and MCs in S1 L5. How these cells develop sharply distinct morphologies given overlapping transcriptomic and electrophysiological profiles, is an interesting open question.

Even though we did not identify any NMCs in V1, the transcriptomic reference dataset ^24^ contained many V1 cells from the *Sst Hpse Cbln4* type, and we found that around a third of MCs from V1 had transcriptomic profile mapping to this type. These cells show an electrophysiological profile intermediate between MCs and NMCs, but morphologically correspond to MCs based on our data. We hypothesize that these cells may be “latent NMCs”, present in V1, but failing to develop a NMC morphology due to the nearly complete absence of stellate cells in V1. Tasic et al. ^24^ showed that the majority of transcriptomic inhibitory types are shared between two very different cortical areas (V1 and ALM). Our findings demonstrate that this does not necessarily imply that morphological types are also all shared.

Using Patch-seq, we also performed single-cell RNA-sequencing of a small number of L5 SOM^+^ cells in both S1 and V1. Morphologically, almost all SOM^+^ cells in V1 L5 (except some fast-spiking cells) ^28,45^ and the majority of SOM^+^ cells in S1 L5 ^21,28,45,61^ are known to be MCs. We found that L5 SOM^+^ cells had electrophysiological features similar to L4 MCs (Fig. 4d), but mostly mapped to a different set of transcriptomic clusters than the L4 SOM^+^ cells (Fig. 4b). These results identify five transcriptomic clusters from Tasic et al. ^24^ as L5 MCs, but the differences between these clusters remain unclear.

### Towards multimodal cell type definition

In this work we have focused on morphologically defined cell types. At the same time, there is a growing understanding that cell type definitions should take into account multimodal information, such as morphology, electrophysiology, and transcriptomics, as opposed to being based on a single modality ^62^. In our V1 L4 data set, we identified seven morphological types of interneurons but only four electrophysiological types (Fig. 2): four PV^+^ could not be distinguished on the basis of their firing as they were all fast-spiking. This is in a qualitative agreement with the findings of Gowens et al. ^60^ who identified twice as many morphological types (m-types) as electrophysiological ones (e-types). We only obtained the transcriptomic information for SOM^+^ neurons, but found out that MCs in V1 L4 could belong to two different transcriptomics types (t-types), one of which corresponded to NMCs in S1; inside V1, the cells from these two t-types had slightly different electrophysiology (Fig. 4). An integrative definition of cell type in V1 should take this into account.

Importantly, the SOM-Cre line does not label neurons in exact correspondence with transcriptomic classes. We found that ∼8% of V1 L4 neurons labeled in the SOM-Cre line were fast-spiking cells with the morphology of basket cells (Fig. 3), in agreement with previous reports ^28,41,42^. In our patch-seq experiments we found three sequenced SOM-Cre^+^ neurons that were fast-spiking and mapped to *Pvalb* transcriptomic types. We did not detect SOM (zero read count) in either of these three cells, suggesting that they likely had transiently expressed it during development, as hypothesized by Hu et al. ^41^ Interestingly, all three cells mapped to the same *Pvalb* type: *Pvalb Reln Itm2a*.

### Circuit organization in L4: V1 vs. S1

In terms of connectivity, both MCs in V1 and NMCs in S1 avoid connecting to each other (apart from forming gap junctions; Fig. S8), and project to excitatory population in L4. Moreover, the axonal morphologies of these two cell types seemed to match the respective dendritic morphologies of their excitatory neuronal targets. In V1, axons of L4 MCs primarily projected to L1 where they are potentially able to synapse onto the tuft of L4 PYRs, similar to the pattern described in other cortical layers ^63,64^. In S1, by contrast, axons of L4 NMCs were more localized, matching the more compact dendritic structure of stellate cells. This observation is in line with previous findings that the excitatory identity controls the survival and wiring of local interneurons ^65,66^. We suggest that the difference in the morphology of SOM^+^ neurons between these two cortical areas might be a result of the difference in dendritic organization of the targeted excitatory neurons. Consistent with this, in S1 L5 where the principal excitatory cells are pyramidal, their inhibitory input comes from L5 MCs ^45^. We hypothesize that the reshaping of excitatory neurons’ apical dendrites in S1 L4 during development, which depends on the sensory input ^7^, could be followed by the corresponding reshaping of SOM^+^ neurons. It will be interesting to test whether this MC/pyramidal and NMC/stellate paring exists in other cortical areas and other species.

On the other hand, while we found that SOM^+^ cells receive inputs from local excitatory neurons in S1 L4, in agreement with previous studies ^14,23^, we did not detect connections from L4 PYRs to L4 MCs in V1. SOM^+^ MCs in other layers are known to receive facilitating excitatory inputs from local principal neurons in both S1 ^67,68^ and V1 ^28,69^. However, our results suggest that L4 MCs in V1 behave differently. Interestingly, previous studies have also shown that in V1, L4 MCs also receive weak inputs from LGN compared to other interneuron types ^20,21^. Within S1, Naka et al. ^45^ showed that L4 excitatory neurons connect to NMCs in L5 but not to MCs in L5, which together with our findings, suggests that even across layers, stellate cells do not target MCs but only NMCs. Further investigations are needed to test whether L4 MCs in V1 are driven by PYRs in other layers or by long-range inputs from other areas.

In addition to examining the connectivity among PYRs and MCs in V1 L4, we tested the connectivity of BCs, another major cell type in L4 (Fig. S10). We found that BCs in L4 followed the same connectivity rules as described for basket cells in other layers ^28^ and in younger animals ^48^: BCs inhibit other BCs, MCs, and pyramidal cells, and are inhibited by MCs and excited by PYRs. All of these connection patterns have also been reported in S1 L4 in young mice ^23^, suggesting that the circuitry wiring involving PV^+^ cells is roughly conserved between these two areas and across age.

Finally, we found very low connection probability between PYRs in V1 L4, which was consistent with the findings in V1 L2/3 and V1 L5 of adult mice ^28^, but much lower than what was reported in young animals ^70,71^. We directly showed that this difference in connection probability among excitatory neurons is due to the age of the animal (Fig. S11).

### Summary

In conclusion, we confirmed the difference in morphology of L4 principal cells and revealed a difference in morphology of L4 SOM^+^ interneurons in V1 and S1 of adult mice. In each area, the morphology of SOM^+^ interneurons matched that of the excitatory neurons, suggesting that one of them might adapt to another. Furthermore, we found differences in the connections from excitatory neurons to SOM^+^ interneurons, suggesting a different functional role of SOM^+^ interneurons in different cortical areas. In addition, we found that there is no one-to-one match between the morphological and the transcriptomic types of SOM^+^ interneurons, highlighting the need of multi-modal profiling of cell types in the neocortex. Our results support the view that cell-type-specific circuit motifs, such as the Martinotti/pyramidal and non-Martinotti/stellate pairs, are used as building blocks to assemble the neocortex.

## Methods

### Data and code availability

Patch-seq data will be made available at https://www.ncbi.nlm.nih.gov/geo/. Apart from the raw reads, it will include a table of read counts, a table of RPKM values, and a table of the extracted electrophysiological features. Morphological reconstructions will be made available at http://neromorpho.org. Raw electrophysiological recordings will be made available at http://zenodo.org.

The analysis code in Python will be made available at http://github.com/berenslab/layer4. This includes data analysis of electrophysiological recordings, data analysis of the morphological reconstructions, and data analysis of the transcriptomic data. This repository also includes a table of the extracted electrophysiological features for the morphological data set.

### Animals

Experiments on adult male and female mice (median age 72, interquartile range 63--88, full range 50--330 days, Fig. S1) were performed using wild-type (*n*=24), Viaat-Cre/Ai9 (*n*=47), Scnna1-Cre/Ai9 (*n*=5 for V1 and *n*=5 for S1), SOM-Cre/Ai9 (*n*=14 for V1 and *n*=19 for S1), VIP-Cre/Ai9 (*n*=8), and PV-Cre/Ai9 mice (*n*=31). Crossing Viaat-Cre mice (Viaat encodes a transporter required for loading GABA and glycine) with Ai9 reporter mice globally labels GABAergic interneurons with the fluorescence marker tdTomato ^37^. SOM-Cre/Ai9 mice, VIP-Cre/Ai9 mice, and PV-Cre/Ai9 mice have SOM^+^ interneurons, PV^+^ interneurons and VIP^+^ interneurons labeled with the fluorescent marker tdTomato respectively. Scnn1a-Cre/Ai9 mice have excitatory neurons in L4 selectively labeled with tdTomato. Additional younger Scnn1a-Cre/Ai9 mice (P15-20, *n*=5; P30-40, *n*=5) were used to study connectivity between excitatory neurons at the different ages. Additional Scnn1a-Cre/Ai9 mice (*n*=8) were used for measuring within-barrel connectivity between excitatory neurons in S1. Additional SOM-Cre/Ai9 mice (*n*=6) were used for patch-seq experiments. Animal preparation procedures for animals maintenance and surgeries were performed according to protocols approved by the Institutional Animal Care and Use Committee (IACUC) of Baylor College of Medicine.

Viatt-Cre line was generously provided by the Dr. Huda Zoghbi’s laboratory. The other Cre lines were purchased from Jackson Laboratory:

- SOM-Cre: http://jaxmice.jax.org/strain/013044.html;
- VIP-Cre: http://jaxmice.jax.org/strain/010908.html;
- PV-Cre: http://jaxmice.jax.org/strain/008069.html;
- Scnn1a-Cre: https://www.jax.org/strain/013044;
- Ai9 reporter: http://jaxmice.jax.org/strain/007909.html.

### Slice preparation

Slice preparation followed methods previously described in Jiang et al. (2015). Briefly, animals were deeply anesthetized using 3% isoflurane. After decapitation, the brain was removed and placed into cold (0−4 °C) oxygenated NMDG solution containing 93 mM NMDG, 93 mM HCl, 2.5 mM KCl, 1.2 mM NaH2PO4, 30 mM NaHCO3, 20 mM HEPES, 25 mM glucose, 5 mM sodium ascorbate, 2 mM Thiourea, 3 mM sodium pyruvate, 10mM MgSO4 and 0.5 mM CaCl2, pH 7.35 (all from SIGMA-ALDRICH). 300 µm thick parasagittal slices were cut and special care was taken to select only slices that had a cutting plane parallel to the apical dendrites to ensure preservation of both axonal and dendritic arborization structures. The slices were incubated at 37.0±0.5 °C in oxygenated NMDG solution for 10-15 minutes before being transferred to the artificial cerebrospinal fluid solution (ACSF) containing: 125 mM NaCl, 2.5 mM KCl, 1.25 mM NaH2PO4, 25 mM NaHCO3, 1 mM MgCl2, 25 mM glucose and 2 mM CaCl2, pH 7.4 (all from SIGMA-ALDRICH) for about 1 h. During recordings, slices were continuously perfused with oxygenated physiological solution throughout the recording session.

### Electrophysiological recordings

Recordings were performed using patch recording pipettes (5−8 MΩ) filled with intracellular solution containing 120 mM potassium gluconate, 10 mM HEPES, 4 mM KCl, 4 mM MgATP, 0.3 mM Na3GTP, 10 mM sodium phosphocreatine and 0.5% biocytin,pH 7.25 (all from SIGMA-ALDRICH). We used two Quadro EPC 10 amplifiers that allowed us to perform simultaneous recordings up to 8 cells. The PatchMaster software and custom-written Matlab-based programs were used to operate the Quadro EPC 10 amplifiers and perform online and offline analysis of the data. In order to extract information about passive membrane properties and firing patterns, neurons’ responses were recorded upon 600 ms long current pulse injections starting from −100 / −200 pA with 20 pA step.

To identify synaptic connections, current pulses were injected into the presynaptic neurons (2 nA for 2 ms at 0.01–0.1 Hz) to evoke AP while post-synaptic membrane potential of other simultaneously recorded neurons were monitored to detect unitary inhibitory or excitatory postsynaptic potentials (uI(E)PSPs). The uIPSPs were measured while the membrane potentials of the putative postsynaptic cells were held at −60±3 mV, whereas uEPSPs were measured while membrane potentials of the putative postsynaptic cells were held at −70±3 mV. Paired-pulse ratio (PPR) was calculated as the ratio between the mean amplitude of the second and the first uEPSC obtained by injecting the presynaptic neuron with two consecutive stimuli of 2nA with 100ms interval. We recorded 10-30 individual traces, average of which was used to calculate uI(E)PSPs amplitude.

Neurons were assigned to L4 based on the neocortical layer boundaries and the small neuronal somata that characterize this layer, which were clearly visible in the micrograph under the bright-field microscope. The layer identity of each neuron was also confirmed *post-hoc* by the visualization of their position after the staining.

Because the synaptic connectivity strongly depends on the inter-soma distance ^28^, we took special care to record from groups of neurons with inter-soma distances less than 250 µm. To make sure that the identified connections were monosynaptic, we morphologically confirmed *post-hoc* the presynaptic neurons for all connections and made sure that the morphology and electrophysiology of the presynaptic neuron for each connection (i.e. pyramidal neurons vs. interneurons) matched the nature of connections (i.e. EPSP vs. IPSP). Indeed, the recovered morphology (i.e. pyramidal neurons vs. interneurons) and EPSP vs. IPSP always matched. Typical recording depth was 15--60 µm, similar to previous studies ^28,70,72^.

Importantly, neuronal structures can be severed (a limitation of all slice electrophysiology experiments) due to the slicing procedure, introducing a potential underestimation of the neuronal morphology and connectivity. However, this did not seem to strongly influence the study of local circuits in the past ^28,73^.

### Staining and morphology recovery

After the end of the patch-clamp recording, the slices were fixed by immersion in freshly-prepared 2.5% glutaraldehyde (from Electron Microscopy Science Cat.no. 16220), 4% paraformaldehyde (from SIGMA-ALDRICH Cat.no. P6148) in 0.1 M phosphate-buffered saline at 4°C for at least 72h. The slices were subsequently processed with the avidin-biotin-peroxidase method in order to reveal the morphology of the neurons. To increase the success rate in recovering the morphology of GABAergic interneurons, especially detail of their fine axonal arbors, we made additional modifications described as previously ^28,29^. The morphologically recovered cells were examined and reconstructed using a 100X oil-immersion objective lens and a camera lucida system. Tissue shrinkage due to the fixation procedure was not compensated for. The shrinkage of the tissue surrounding the biocytin-stained cells was about 10--20%, consistent with previous studies ^28,50^.

For barrel identification and comparison with tdTomato signal in Scnn1a-Cre mice, we performed cytochrome C staining following protocols described in the literature ^3,74^. To find the barrel locations in images with cytochrome C and tdTomato (Fig. S10A), we averaged the pixel intensities as a function of horizontal position within the L4. The resulting intensity trace was normalized to lie between 0 and 1 and high-pass filtered to compensate for the uneven brightness of the images. To do the high-pass filter, we used a Fourier function of the form:

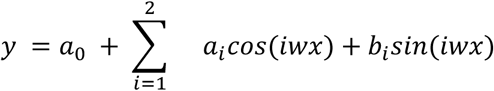

that was fitted to each trace (*w* was fitted along the *a*_i_ and *b*_i_ coefficients) and then subtracted from it. The signal from the cytochrome C was inverted to match the directionality of the tdTomato signal. Barrel center locations were estimated as the positions of the peaks after smoothing with a σ=250 µm Gaussian filter.

### Patch-seq procedure and sequencing

To obtain electrophysiology and transcriptome data from single neurons, we used our recently described Patch-seq protocol ^29^ with the following additional modifications. Recording pipettes of 5 MΩ resistance were filled with RNase-free intracellular solution containing: 101 mM potassium gluconate, 4 mM KCl, 10 mM HEPES, 0.2 mM EGTA, 4 mM MgATP, 0.3 mM Na_3_GTP, 5 mM sodium phosphocreatine (all from SIGMA-ALDRICH), and 1 U/μl recombinant RNase inhibitor (Takara Cat.no. 2313A), pH ∼7.25. The cDNA was amplified using 18 amplification cycles and the size distribution and concentration of the libraries were analyzed with an Agilent Bioanalyzer 2100. cDNA samples containing less than 1.5 ng total cDNA, or with an average size less than 1,500 bp were not sequenced.

To construct the final sequencing libraries, 0.2 ng of purified cDNA from each sample was tagmented using the Illumina Nextera XT Library Preparation using ⅕ of the volumes stated in the manufacturer’s recommendation. The DNA was sequenced from single end (75 bp) with standard Illumina Nextera i5 and i7 index primers (8 bp each) using an Illumina NextSeq500 instrument. Investigators were blinded to cell type during library construction and sequencing.

Reads were aligned to the mouse genome (`mm10` assembly) using STAR (v2.4.2a) with default settings. We only used read counts (and not RPKM values, number of reads per kilobase of transcript per million total reads) for all data analysis presented here, but for completeness we mention that RPKM values were computed using `rpkmforgenes` ^75^ and NCBI RefSeq gene and transcript models (downloaded on the 24th of June 2014).

### Data analysis of the morphological reconstructions

Reconstructed morphologies of *n*=92 cells were converted into SWC format and further analyzed using custom Python code (see the github repository linked above). Each cell was soma-centered and rotated such that the *z* coordinate (height) was oriented along the cortical depth and the *x* coordinate (width) was oriented along the first principal component of the *xy* point cloud, i.e. roughly corresponded to the cell’s largest extent in the plane orthogonal to the cortical depth. For further analysis we computed and combined two different feature representations of each cell: the XZ density map and a set of morphometric statistics.

#### XZ density map

We sampled equidistant points with 100 nm spacing along each neurite and normalized the resulting point cloud such that the smallest coordinate across all points of all cells was 0 and the largest was 1 ^76^. The normalized point cloud was projected onto the *xz*-plane and binned into 100× 100 bins spanning [-0.1, 1.1]. We smoothed the resulting density map by convolving it with a 11× 11 bin Gaussian kernel with standard deviation *σ*=2. For the purposes of downstream analysis, we treated this as set of 10,000 features.

#### Morphometric statistics

For each cell we computed a set of 16 summary statistics: number of branch points, cell width, cell depth, cell height, number of tips, number of stems, total neurite length, maximal neurite length, maximum branch order, maximal segment length, average tortuosity, maximal tortuosity, average branch angle, maximal branch angle, average path angle, and maximal path angle.

#### Pairwise classification

We followed the pipeline that we recently benchmarked in Laturnus et al. ^43^. As predictors for pairwise classification we used morphometric statistics and density maps. Due to the very high dimensionality of the density maps, we reduced them to 10 principal components (for cross-validation, PCA was computed on each outer-loop training set separately, and the same transformation was applied to the corresponding outer-loop test set). This makes the final feature dimensionality equal to 36.

For classification, we used logistic regression regularized with elastic net. Regularization parameter alpha was fixed to 0.5, which is giving equal weights to the lasso and ridge penalties. We used nested cross-validation to choose the optimal value of the regularization parameter lambda and to obtain an unbiased estimate of the performance. The inner loop was performed using the `civisanalytics` Python wrapper around the `glmnet` library ^77^ that does K-fold cross-validation internally. We used 5 folds for the inner loop. We kept the default setting which uses the maximal value of lambda with cross-validated loss within one standard error of the lowest loss (`lambda_best`) to make the test-set predictions:

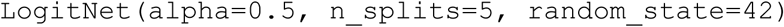

Note that the default behavior of `glmnet` is to standardize all predictors. The outer loop was 10 times repeated stratified 5-fold cross-validation, as implemented in `scikit-learn` by

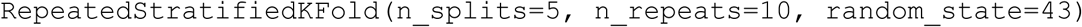

Outer-loop performance was assessed via test-set accuracy.

#### t-SNE

For the t-SNE visualization, we reduced density maps and morphometric statistics of the *n*=92 cells to 10 principal components each. We scaled each set of 10 PCs by the standard deviation of the respective PC1, to make three sets be roughly on the same scale. Then we stacked them together to obtain a 20-dimensional representation of each cell. Exact (non-approximate) t-SNE was run with perplexity 15, random initialisation with seed 42, and early exaggeration 4, using `scikit-learn` implementation:

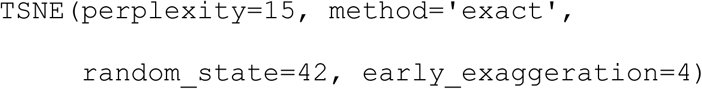

### Automatic extraction of electrophysiological features

Thirteen electrophysiological features were automatically extracted using Python scripts from the Allen Software Development Kit (https://github.com/AllenInstitute/AllenSDK) with additional modifications to account for our experimental paradigms (see the github repository linked above). An illustration of the feature extraction procedure for one exemplary neuron is shown in Fig. S1. Here we briefly specify how each feature was extracted.

The resting membrane potential and the input resistance were computed differently for the standard patch-clamp/morphology recordings and for the Patch-seq recordings, because of the differences in the stimulation protocol between these two sets of experiments. In the Patch-seq experiments, the current clamp value before each current stimulation was fixed at 0 pA for all cells. Consequently, we computed the resting membrane potential as the median membrane voltage before stimulation onset. Input resistance for each hyperpolarizing stimulation was calculated as the ratio of the maximum voltage deflection to the corresponding current value. We took the median of all hyperpolarizing currents as the final input resistance value. In contrast, in the standard patch-clamp experiments, the current clamp before current stimulation was not always fixed at 0 pA. For that reason we used linear regression (for robustness, random sample consensus regression, as implemented in `scikit-learn`) of the steady state membrane voltage onto the current stimulation value to compute the input resistance (regression slope) and the resting membrane potential (regression intercept) (Fig. S1D). For this we used five highest hyperpolarizing currents (if there were fewer than five, we used those available).

To estimate the rheobase (minimum current needed to elicit any spikes), we used robust regression of the spiking frequency onto the stimulation current using the five lowest depolarizing stimulation currents with non-zero spike count (if there fewer than five, we used those available) (Fig. S1D). The point where the regression line crosses the *x*-axis gives the rheobase estimate. We restricted the rheobase estimate to be between the highest current clamp value eliciting no spikes and the lowest current clamp value eliciting at least one spike. In the rare cases when the regression line crossed the *x*-axis outside of this interval, the nearest edge of the interval was taken instead as the rheobase estimate.

The action potential (AP) threshold, AP amplitude, AP width, afterhyperpolarization (AHP), afterdepolarization (ADP), and the first spike latency were computed as illustrated in Fig. S1C, using the very first spike fired by the neuron. AP width was computed at the half AP height.

The adaptation index (AI) is defined as the ratio of the second interspike interval to the first one (Fig. S1B). We took the median over the five lowest depolarizing current stimulation that elicited at least three spikes (if fewer than five were available, we used all of them).

The maximum number of spikes simply refers to the maximum number of spikes emitted in the 600 ms stimulation window over all stimulation currents (Fig. S1A). The membrane time constant (tau) was computed as the time constant of the exponential fit to the first phase of hyperpolarization (median over all hyperpolarizing traces). Finally, the sag ratio is defined as the ratio of the maximum membrane voltage deflection to the steady state membrane voltage during the first (the lowest) hyperpolarizing current clamp stimulation.

### Data analysis of the electrophysiological features

For the t-SNE visualization (Fig. 2B), we log-transformed the AI values because this feature had a strongly right-skewed distribution (Fig. S1). We also excluded ADP and latency; ADP because it was equal to zero for most neurons and rare cells with non-zero values appeared as isolated subpopulations in the t-SNE representation, and latency because it had high outliers among the FS types, also yielding isolated subpopulations. The remaining 11 features were *z*-scored and exact (non-approximate) t-SNE was run with perplexity 15 and random initialisation with seed 42 using `scikit-learn` implementation:

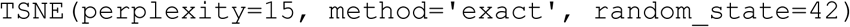

For pairwise classification, we used exactly the same procedure as described above for pairwise classification using the reconstructed morphologies (nested cross-validation with logistic regression regularized with elastic net). All 13 features were used, with log-transformed AI and log-transformed latency (as shown in Fig. S1).

### Data analysis of the RNA-seq data

#### Quality control

The total number of sequenced cells was *n*=118. Four cells were excluded because the sum of counts across all genes (library size) was below 1500 (Fig. S3a). The remaining *n*=114 cells were mapped to the full set of 133 transcriptomic clusters identified in Tasic et al. ^24^; see below for the details. One cell was excluded because it mapped to one of the excitatory types, and three cells were excluded because they mapped to *Pvalb Reln Itm2a* type (and were fast-spiking). All the remaining *n*=110 cells mapped to the *Sst* clusters. Among those, eight cells did not have good electrophysiological recordings (the recordings were either lost or were of bad quality) and were excluded from all downstream analyses that required electrophysiological data (leaving *n*=102 cells).

#### Mapping to the reference clusters

Using the count matrix of Tasic et al. (*n*=23,822, *d*=45,768), we selected 3000 “most variable” genes (see below). We then log-transformed all counts with log_2_(x+1) transformation and averaged the log-transformed counts across all cells in each of the 133 clusters, to obtain reference transcriptomic profiles of each cluster (133×3000 matrix). Out of these 3000 genes, 2686 were present in the `mm10` reference genome that we used to align reads in our data (see above). We applied the same log_2_(x+1) transformation to the read counts of our cells, and for each cell computed Pearson correlation across the 2686 genes with all 133 Tasic et al. clusters. Each cell was assigned to the cluster to which it had the highest correlation.

#### Gene selection

To select “most variable” genes, we found genes that had, at the same time, high non-zero expression and high probability of near-zero expression ^78^. Our procedure is described in more detail elsewhere ^47^. Specifically, we excluded all genes that had counts of at least 32 in fewer than 10 cells. For each remaining gene we computed the mean log_2_ count across all counts that were larger than 32 (non-zero expression, *μ*) and the fraction of counts that were smaller than 32 (probability of near-zero expression, *τ*). Across genes, there was a clear inverse relationship between *μ* and *τ*, that roughly follows exponential law *τ ≈* exp*(−1.5·μ+a)* for some horizontal offset *a*. Using a binary search, we found a value *b* of this offset that yielded 3000 genes with *τ >* exp*(−1.5·μ+b) + 0.02*. These 3000 genes were selected.

#### t-SNE

The t-SNE visualization of the whole Tasic et al. ^24^ dataset shown in Fig. S3C was taken from our previous work ^47^. It was computed there using scaled PCA initialization and perplexity combination of 30 and 238 (1% of the sample size), following preprocessing steps of library size normalization (by converting counts to counts per million), feature selection (3000 most variable genes), log_2_(x+1) transformation, and reducing the dimensionality to 50 using PCA.

To make t-SNE visualization of the somatostatin part of the Tasic et al. dataset (Fig. 4b), we selected all cells from all *Sst* clusters apart from the very distinct *Sst Chodl* (20 clusters, 2701 cells). Using these cells, we selected 500 most variable genes using the same procedure as described above. We used the same preprocessing steps as above, perplexity 50, and scaled PCA initialisation ^47^.

#### Mapping to t-SNE

For each of the *n*=110 Patch-seq cells, we computed its Pearson correlation with each of the 2701 reference cells across the 500 genes, most variable in the somatostatin part of the Tasic et al. data set (only 472 genes present in our data were used). Then we found 10 reference cells with the highest correlations (10 “nearest neighbours” of the Patch-seq cell) and positioned our cell at the coordinate-wise median t-SNE location of those 10 reference cells ^47^.

#### Mapping to somatostatin clusters

The mapping of the *n*=110 Patch-seq cells to the 20 somatostatin clusters (Fig. 4C, S3) was done exactly as the mapping to the full set of 133 clusters described above, but this time only using 500 genes, most variable in the somatostatin part of the Tasic et al. ^24^ data set (only 472 genes present in our data were used).

#### Sparse reduced-rank regression

We used our implementation of sparse reduced-rank regression (RRR) described in detail elsewhere ^46^. For the analysis shown in Fig. 4e, we selected 1000 most variable genes as described above, using *n*=102 Patch-seq cells with high-quality electrophysiological recordings. The gene counts were converted to counts per million and log_2_(x+1)-transformed. The columns of the resulting 102×1000 expression matrix were standardized. All electrophysiological features were standardized as well. The rank of RRR was fixed at 2. We used 10-fold cross-validation to select the values of alpha and lambda regularization parameters that would maximize the predicted R-squared. This yielded alpha=0.5 and lambda=1 (with “relaxed elastic net” ^46^). Fig. 4E shows scatter plots of the two standardized RRR components in the transcriptomic and in the electrophysiological spaces. Features and genes are depicted as lines showing correlations of a feature/gene with each of the two components. In the electrophysiological space, all features are shown. In the transcriptomic space, only genes selected by the model are shown. The values of R-squared and correlations between the components from electrophysiological and transcriptomic spaces reported in the caption of Fig. 4e are cross-validation estimates.

## Supplementary text and figures

### Detailed description of interneuron cell types

BCs were the most abundant interneuron type (37.6%, 88/234). Somata of these neurons were usually larger than those of other L4 neurons. Their dendrites projected vertically in a bi-tufted manner, without a complex horizontal structure. The most salient morphological feature of BCs was a thick axon originating from the apical side of the soma. It typically projected towards L2/3 before forming a series of major branches that extensively spread above the apical region of the soma with few branches projecting horizontally and vertically downward to L5. All BCs exhibited a fast-spiking (FS) firing pattern with narrow AP width and high maximal firing rate (Fig. 1b).

MCs (20.1%, 47/234) were similar to those previously described both in developing cortex and in mature cortex in other layers ^28,79–81^. They had bi-tufted dendrites with vertically or obliquely oriented branches. All of them had an ascending axon that projected to L2/3 and L1, where it ramified horizontally and formed a dense axonal cluster of variable extension. A small subset of MCs (8.9%, 11 out of all 124 recovered MCs) showed a secondary axonal cluster within L4 (e.g. the last two MCs in Fig. 1a). Firing pattern and electrophysiological properties showed a strong correspondence to L2/3 and L5 MCs described in both adult ^28^ and developing cortex ^81^. In particular, these neurons were distinguished from other interneurons by their large membrane time constant (Fig. 1b).

BPCs (12.4%, 29/234) had a small soma and bipolar dendrites projecting to L1 and L5. The ascending dendrites formed a tuft in L1, similar to the structure of apical dendrites of PYRs. However, their dendrites lacked dendritic spines. The axon emerged from one of the descending dendrites and projected predominantly to L5. All BPCs showed an irregular-firing pattern associated with a high input resistance and large AP amplitude (Fig. 1b).

NGCs (11.5%, 27/234) were characterized by a very thin axon that highly ramified and formed a dense arborization around cell bodies. These neurons fired late-spiking action potentials with large AP width and high AP threshold (Fig. 1b).

SCs (9.4%, 22/234) were similar to shrub cells that have been previously described in L5 of adult mouse ^28^ and small BCs in L2/3 and L4 of young rats ^28,82^. These neurons had non-polarized dendritic branches mostly residing in L4 and a thick axon often emerging from the apical region of the cell bodies and branching locally around their soma. All SCs exhibited an FS firing pattern (Fig. 1b).

DBCs (5.6%, 13/234) had large cell bodies and vertically-oriented bi-tufted dendrites, similar to BCs. However, unlike BCs, the thick axon emerged often from the bottom of the soma, projecting shortly towards L5 and forming several branches that projected upwards to L2/3 and downwards to L5 with variable distances. Notably, the axons of these cells extended also horizontally into L2/3 and L5, differing slightly from DBCs previously described in L2/3 ^28^. All DBCs exhibited an FS firing pattern (Fig. 1b).

HECs (3.4%, 8/234), with their horizontally extended axonal branches, were similar to the HECs previously reported in L5 ^28,83^. In particular, the axon had a thick primary structure that often emerged from the apical side of the soma and bifurcated into secondary structures that spread horizontally mostly within L4. All HECs exhibited an FS firing pattern (Fig. 1b).

## Supplementary Figures

**Figure S1:**
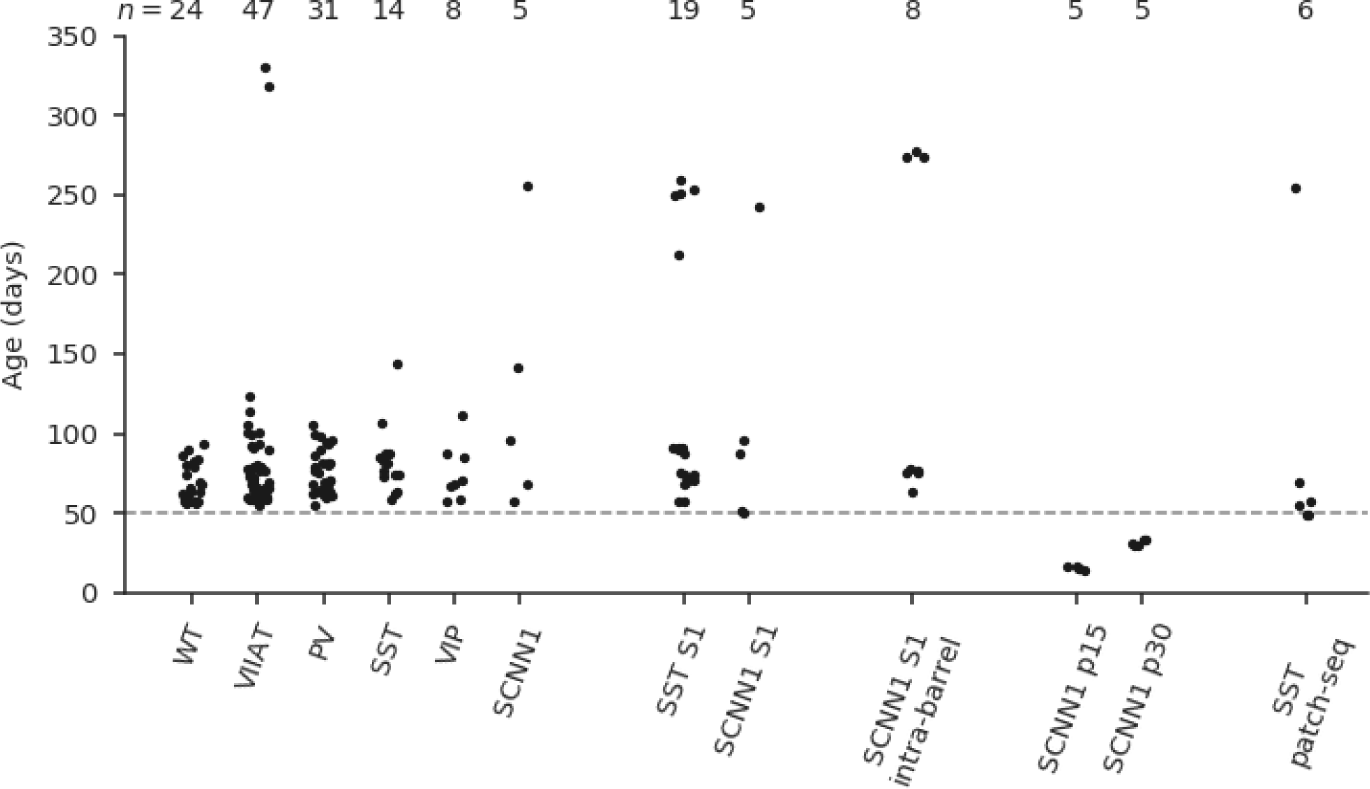
Mice ages. WT stands for wild type, all other abbreviations correspond to Cre lines.

**Figure S2:**
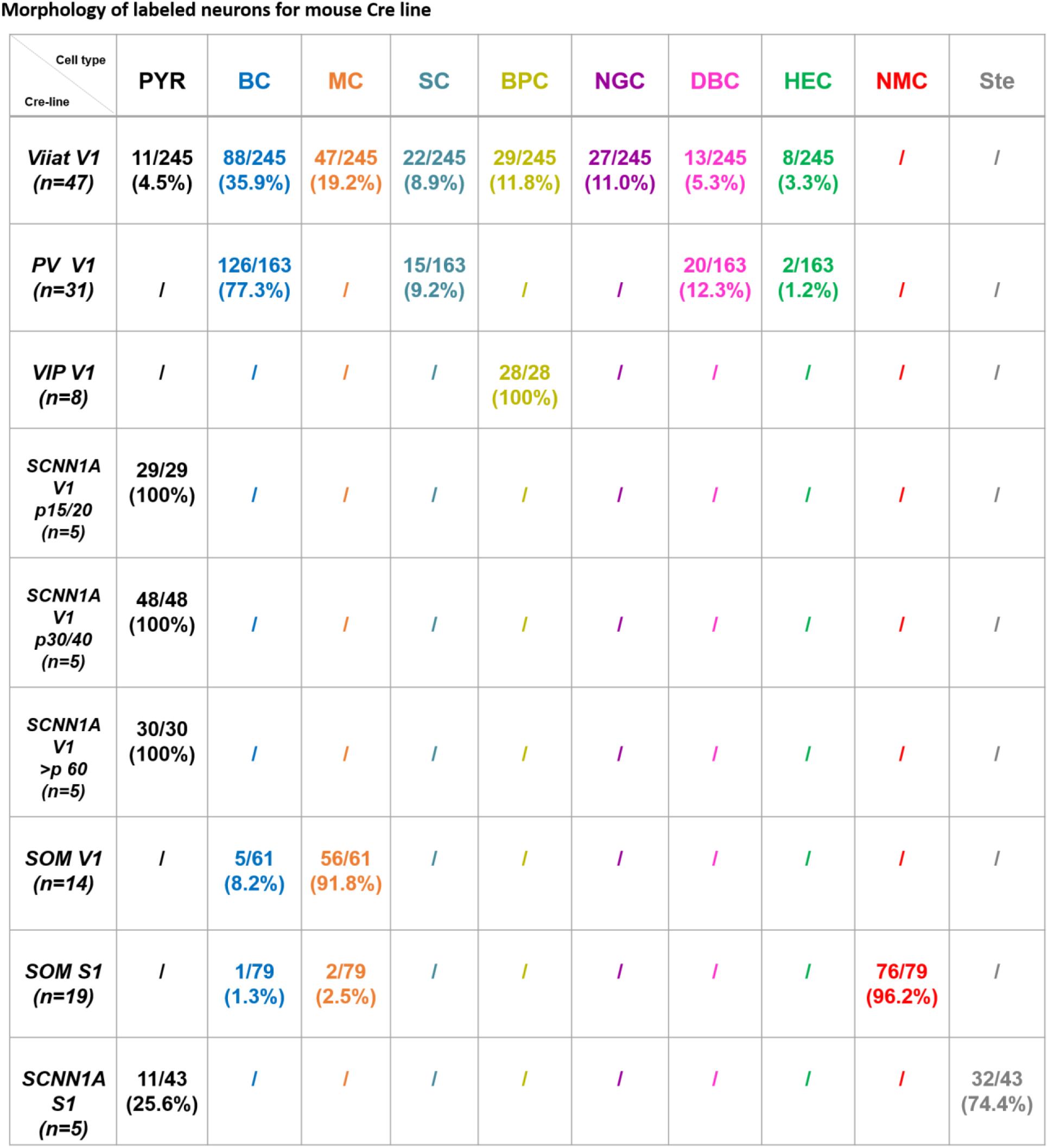
Morphological types of labeled neurons in different mouse Cre lines.

**Figure S3:**
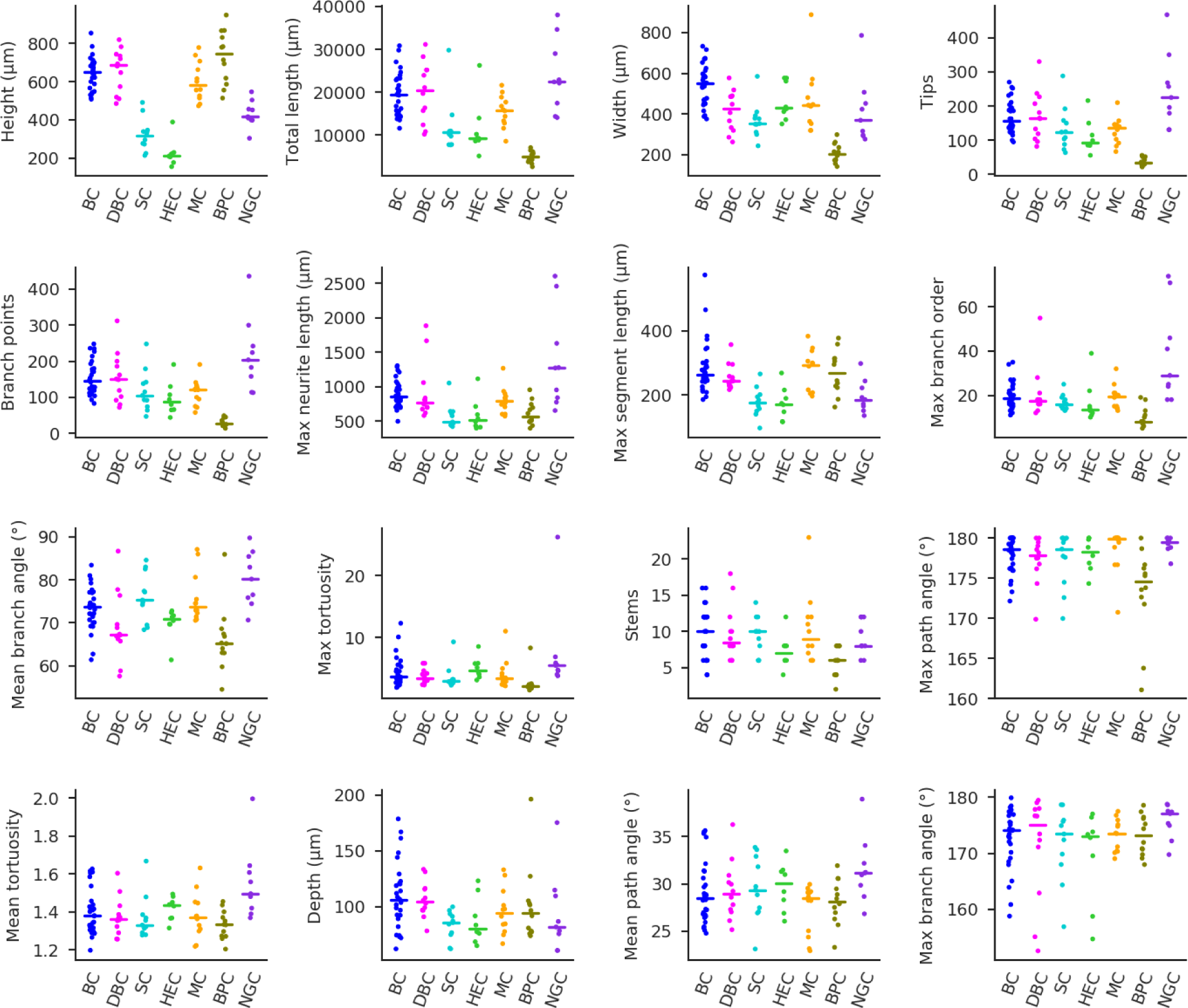
Morphological features of neural cell types in V1 L4. Each panel shows one of the 16 selected morphological summary statistics for *n*=92 neurons: height (extent along the cortical depth), total length of neurites, width (extent along the *x*-axis), number of tips, number of branch points, length of the longest neurite from tip to soma, length of the longest segment, maximum branch order, mean branch angle between two branches, maximum tortuosity, maximum path angle, number of stems extending from the soma, mean tortuosity, depth (extent along the *y*-axis), mean path angle and maximum branch angle. Features are sorted by how strongly they varied between cell types (from strongest to weakest), as quantified by the Kruskal-Wallis test statistic. Horizontal lines show medians in each cell type.

**Figure S4:**
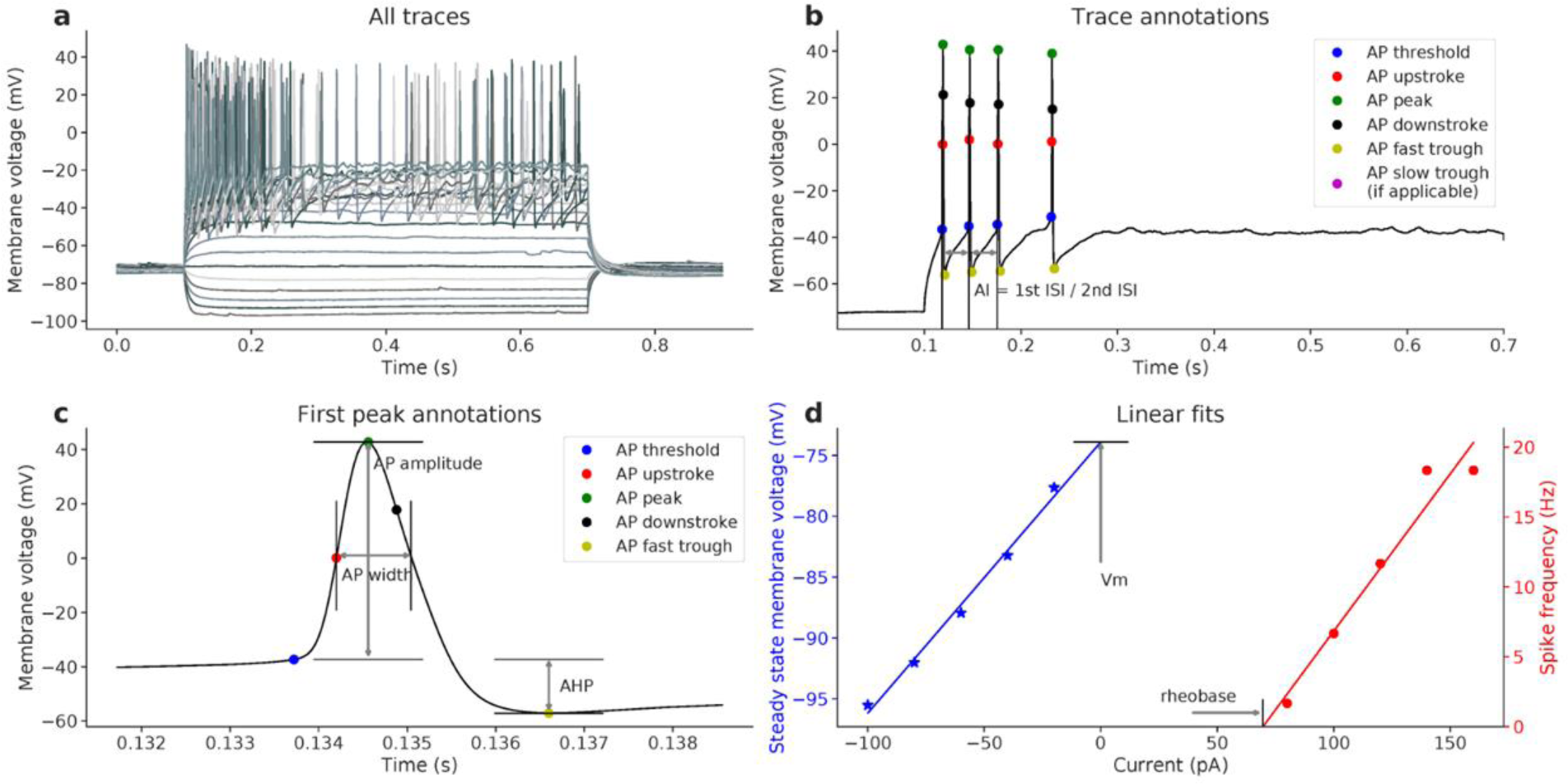
Schematic of the electrophysiological features and the extraction algorithm. All panels show data from the same exemplary Martinotti cell. **(a)** Responses to the consecutive current clamp stimulation currents. The maximum number of spikes elicited in 600 ms was 11. Hyperpolarizing currents are used to compute sag ratio (1.3) and membrane time constant tau (23.2 ms). **(b)** Zoom-in to one particular trace in (a) showing trace annotations and AI (1.13). **(c)** Zoom-in to the first spike elicited by this neuron. This action potential is used to compute AP threshold (−40.1 mV), AP amplitude (71.3 mV), AP width (0.72 ms), AHP (−14.1 mV), ADP (6.6 mV), and latency of the first spike (78.7 ms) **(d)** Blue regression line gives an estimate of resting membrane potential (−58.9 mV) and input resistance (235.8 MΩ). Red regression line gives a rheobase estimate (40 pA).

**Figure S5:**
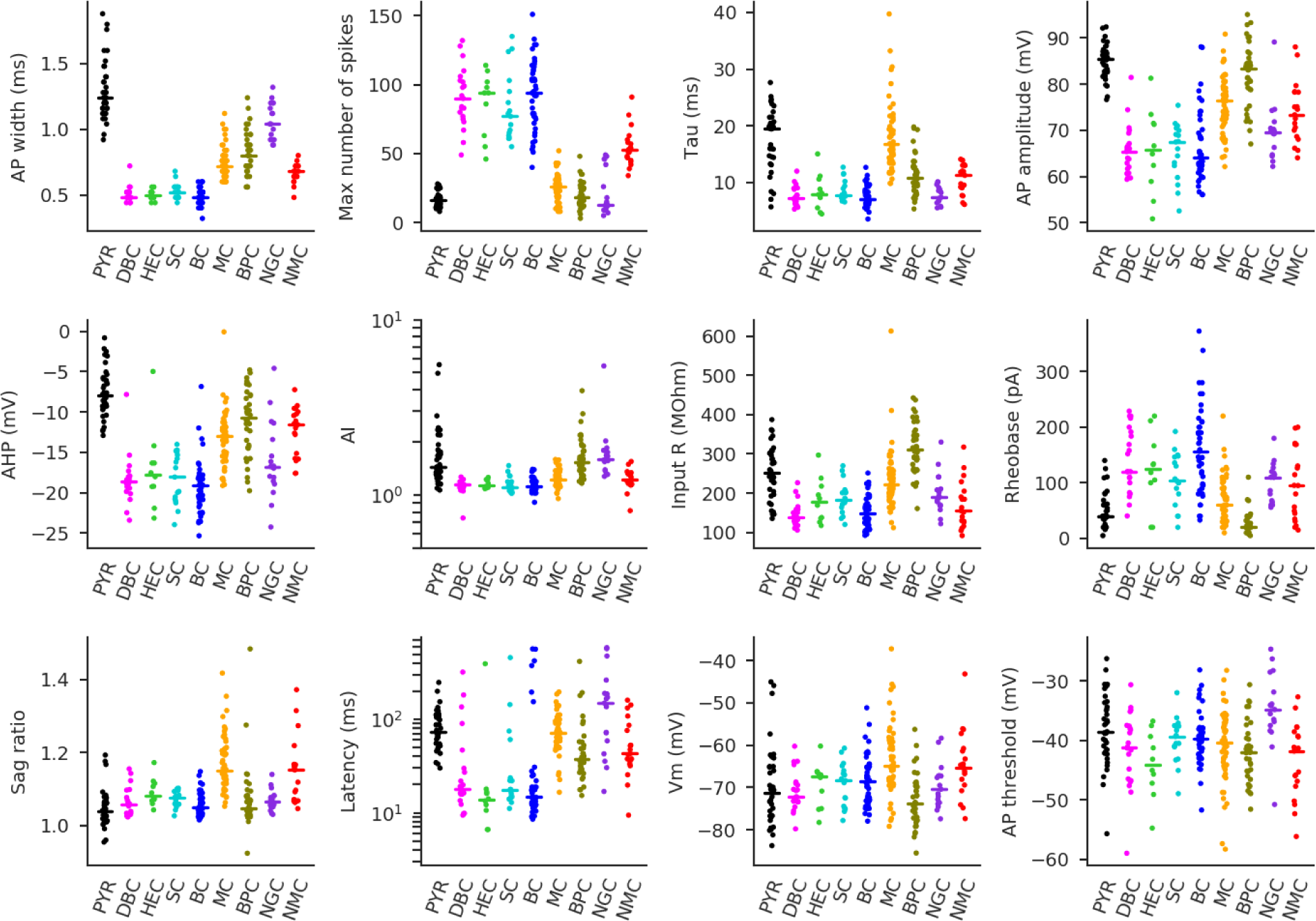
Electrophysiological features of neural cell types. All cell types apart from NMC (red) are from V1 L4. NMC is from S1 L4. Each panel shows one of the 13 automatically extracted electrophysiological features for *n*=254 neurons: action potential (AP) width, maximum number of spikes emitted during 600 ms of stimulation, membrane time constant tau, AP amplitude, afterhyperpolarization (AHP) depth, input resistance, adaptation index, rheobase, sag ratio, latency of the first spike, membrane potential, and AP threshold. Features are sorted by how strongly they varied between cell types (from the most strongly to the least strongly), as quantified by the Kruskal-Wallis test statistic. Horizontal lines show medians in each cell type. Afterdepolarization (ADP) height is not shown because its median was 0 for all cell types. See Fig. S4 for explanations of the features.

**Figure S6:**
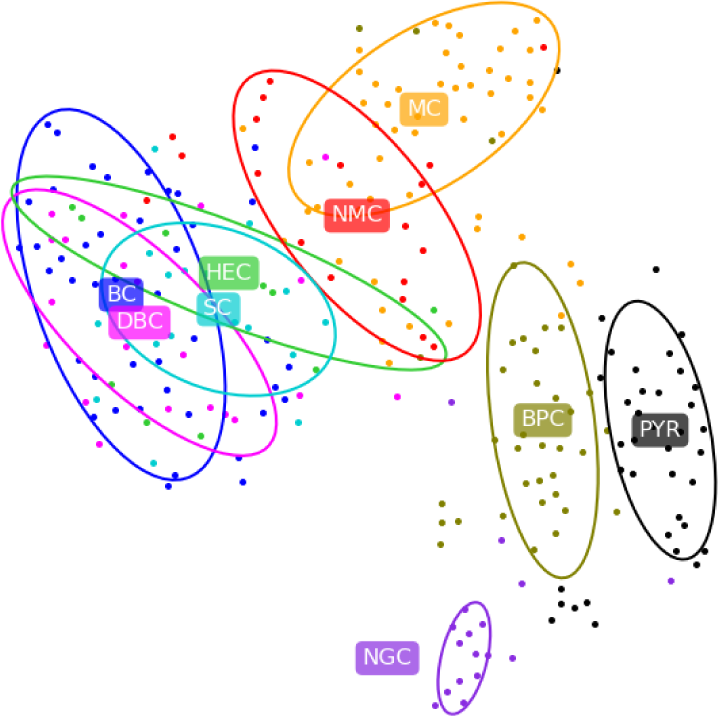
2D visualisation of cell types in the space of electrophysiological features using t-SNE. This figure is analogous to Fig. 2b, but includes *n*=19 NMCs from S1 in addition to the *n*=235 cells from V1.

**Figure S7:**
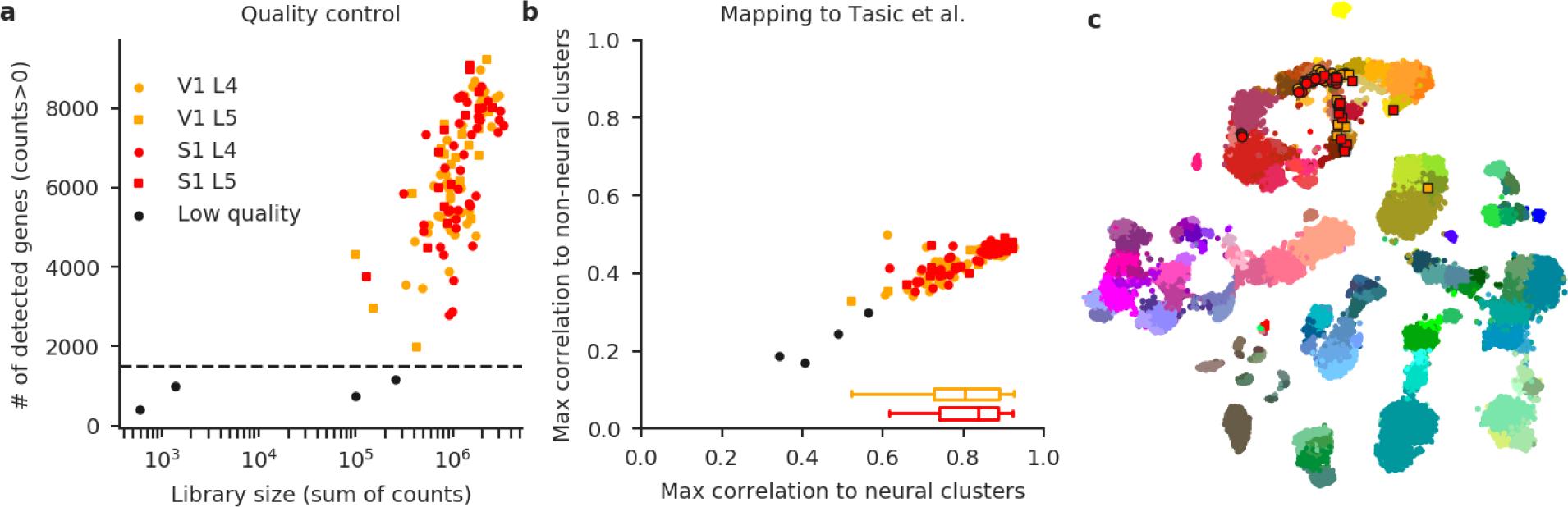
Quality control of Patch-seq data including L5 cells. **(a)** Distribution of library sizes (total sum of gene counts) and numbers of detected genes (number of positive counts) for each sequenced cell (*n*=118). Four cells with less than 1500 genes detected were excluded. **(b)** For each cell, we found its maximal correlation to the cluster means of the Tasic et al. ^24^ dataset across all neural clusters (*x*-axis) and across all non-neural clusters (*y*-axis). Boxplots show distribution of maximal correlation for V1 and S1 cells (excluding the low quality cells). Correlations were not lower for S1 cells, despite the fact that the Tasic et al. dataset only contained data from V1 and ALM. **(c)** All *n*=114 remaining cells were positioned on the t-SNE map of the Tasic et al. dataset. Three cells mapped to Pvalb clusters and one cell mapped to excitatory clusters.

**Figure S8:**
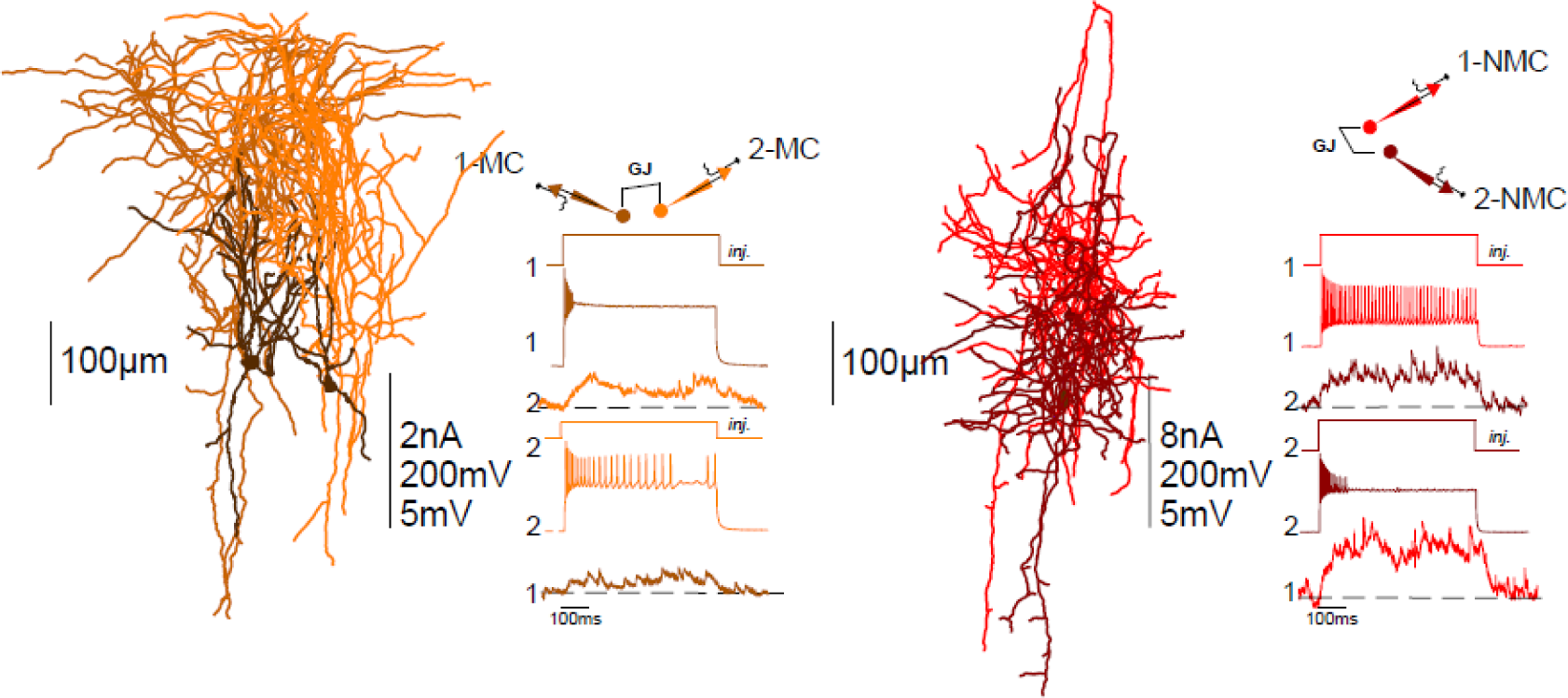
Gap junctions are common between both MCs and NMCs. Schematic representations of simultaneous recordings between L4 MCs in V1 (left) and L4 NMCs in S1 (right). Depolarizing current injections into either MC (left) or NMC (right) were transmitted to the other cell, confirming electrical coupling. The percentage of gap junctions was 23.5% in V1 (8/34) and 30.7% in S1 (8/26).

**Figure S9.**
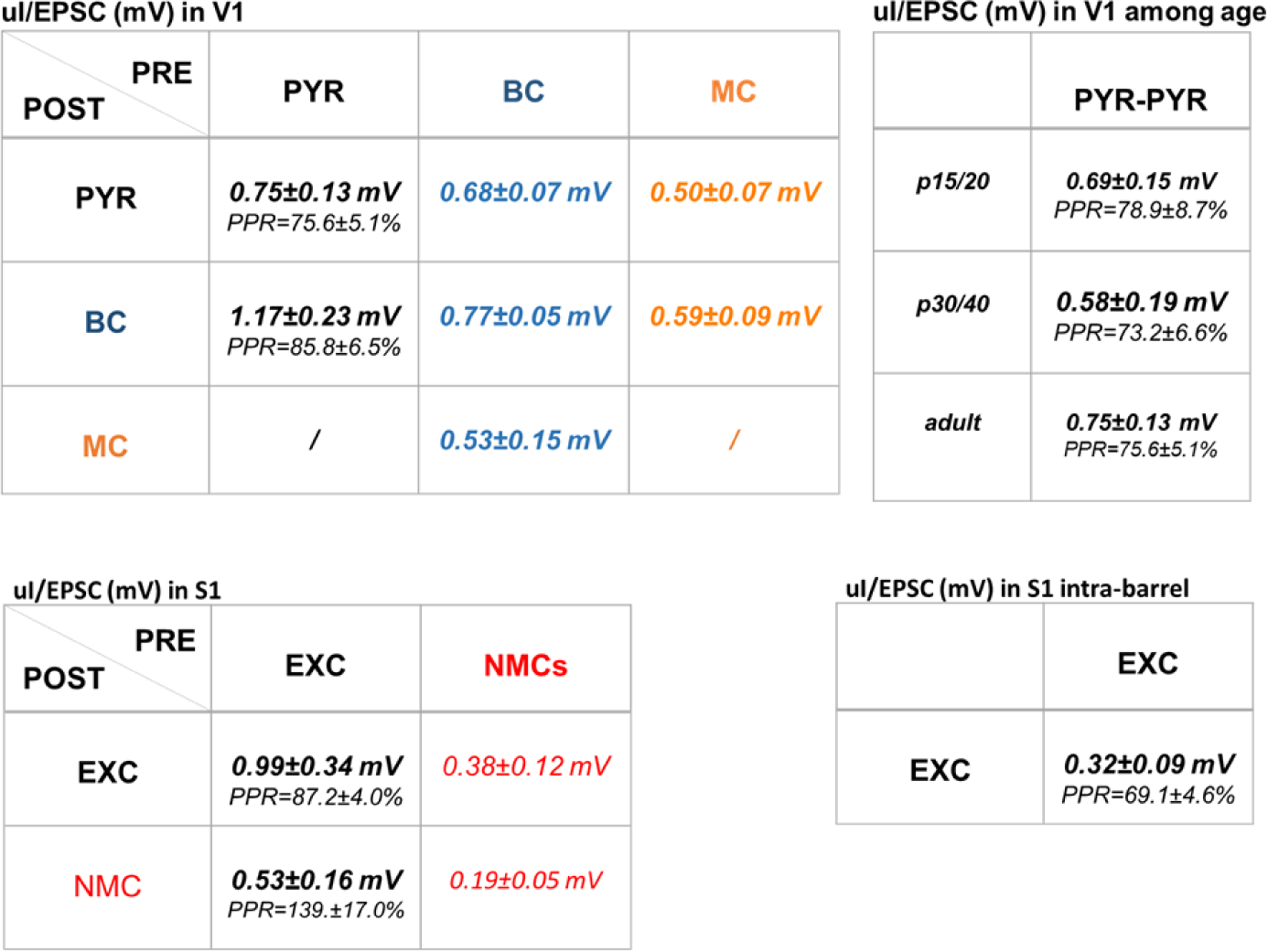
Amplitudes of u(E)psc and (I)PSC.

**Figure S10:**
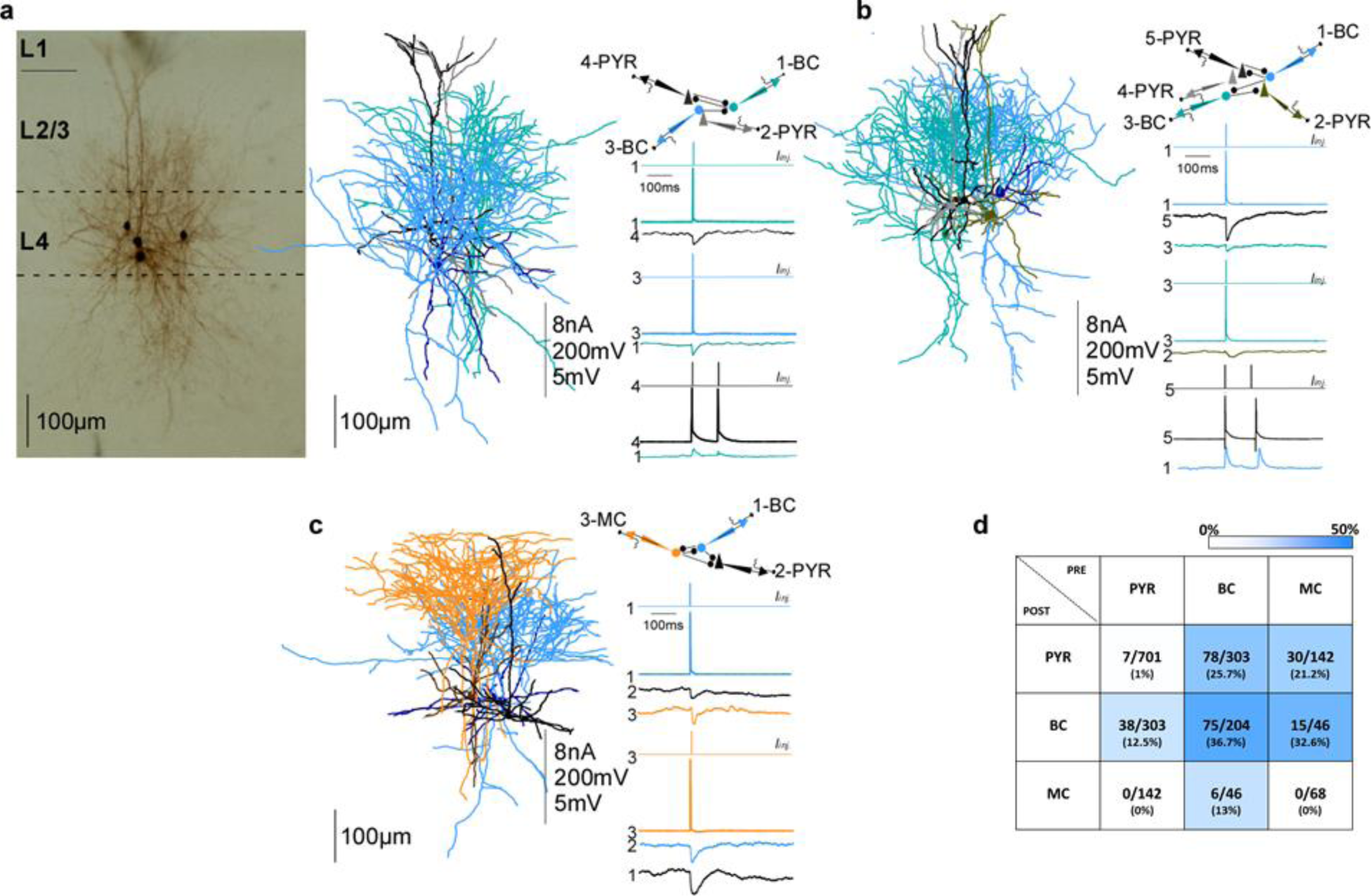
Connectivity between L4 PYRs, BCs, and MCs in V1. **(a)** On the left: Example of morphological recovery of four neurons. Recorded neurons were close to each other (generally less than 250μm). On the right: connection diagram of the same neurons, including two BCs and two PYRs, and their reconstructed morphology. Vertical scale bar indicates: amplitudes of injected current in nA, amplitude of APs in mV and amplitude of uEPSP or uIPSP in mV. **(b)** Connections between five simultaneously recorded neurons including three PYRs and two BCs. **(c)** Connections between three simultaneously recorded neurons including one PYR, one BC and one MC. **(d)** Color coded connectivity matrix showing the connection probabilities between PYRs, BCs and MCs computed as a fraction of all tested connections. Average of uEPSP and uIPSP as well as PPR are reported in Fig. S7.

**Figure S11:**
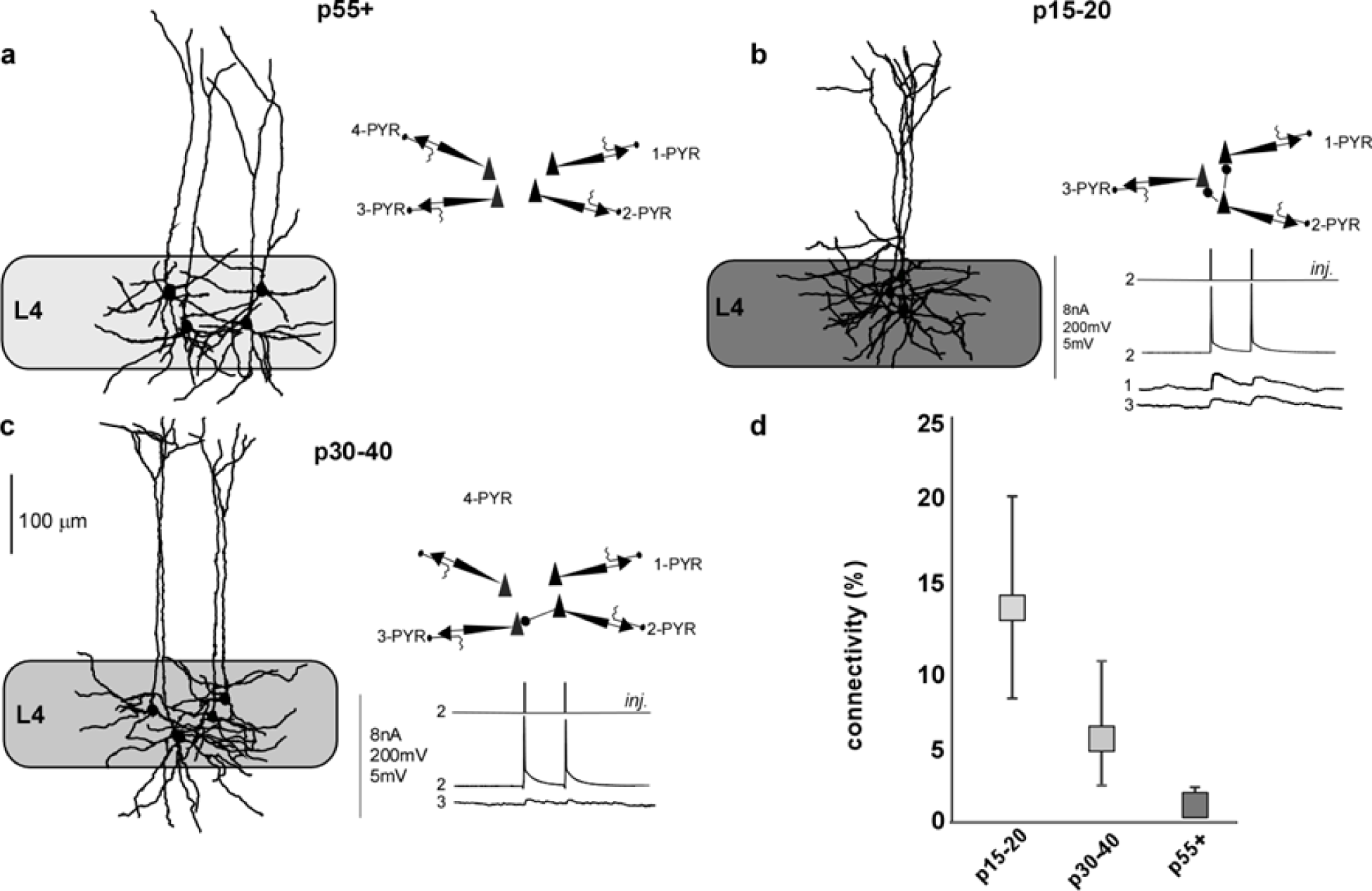
Connectivity between L4 PYRs in V1 at different ages. **(a-c)** Examples of simultaneously recorded L4 neurons in P55+ (A), P15-20 (B), and P30-40 (C) mice. **(d)** Connectivity probability between PYRs at different ages: 13.2% in P15-20 (15/114), 5.1% in P30-40 (8/156), and 1.0% in P55+ with median age p71 (7/701). Error bars are 95% Clopper-Pearson confidence intervals.

**Figure S12:**
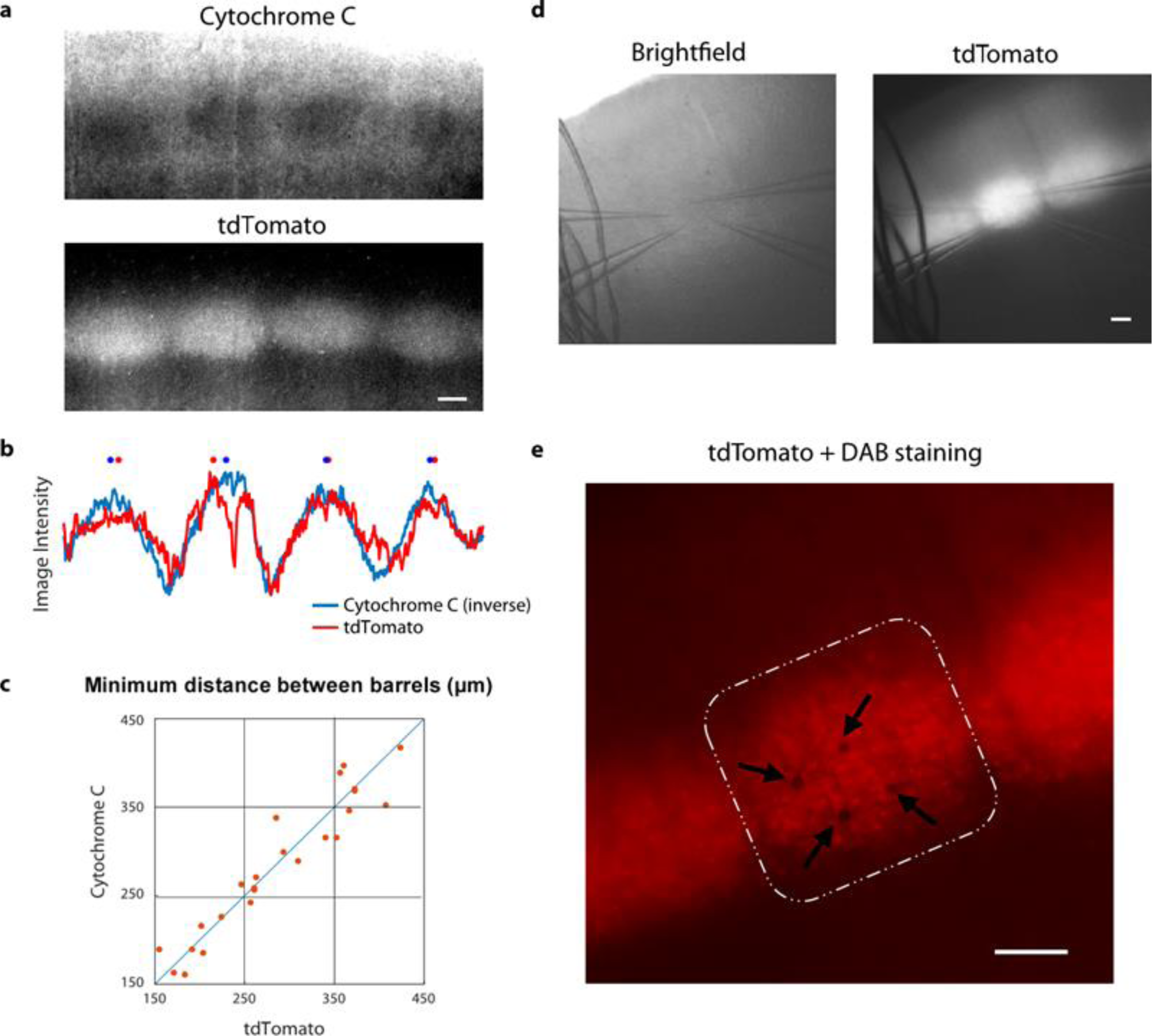
Identification of L4 barrels in S1 for intra-barrel recordings: **(a)** One exemplary slice showing barrels in L4 of somatosensory cortex identified by either cytochrome C staining (above) or by tdTomato fluorescence (below) in Scnn1a-Cre mouse. **(b)** Average intensity for the Cytochrome C and tdTomato images shown in A. Note that the traces shown here are normalized (between 0 and 1) and high-pass filtered but not smoothed. The signal of the cytochrome C was inverted for easier comparison to the fluorescence trace. The barrel centers, indicated with asterisks, were detected as the peaks of intensity of the smoothed traces. **(c)** Distances between adjacent barrels detected with either Cytochrome C or tdTomato. Summary across *n*=7 slices. **(d)** An example of intra-barrel quadruple recording. Left: under brightfield. Right: tdTomato signal. **(e)** Diaminobenzidine (DAB) staining confirmed the intra-barrel localization of the recorded neurons. All scale bars represent 100µm.

## Acknowledgements

We thank Alex Naka for comments and suggestions. Supported by the Intelligence Advanced Research Projects Activity (IARPA) via Department of Interior/Interior Business Center (DoI/IBC) contract number D16PC00003. The U.S. Government is authorized to reproduce and distribute reprints for Governmental purposes notwithstanding any copyright annotation thereon. The views and conclusions contained herein are those of the authors and should not be interpreted as necessarily representing the official policies or endorsements, either expressed or implied, of IARPA, DoI/IBC, or the U.S. Government.

This work was also supported by the German Research Foundation (BE5601/4-1, SFB1233 - 276693517) and the German Excellence Strategy (EXC 2064 - 390727645), the Federal Ministry of Education and Research (FKZ 01GQ1601) and the National Institute of Mental Health and the National Institute of Neurological Disorders And Stroke under Award Number U19MH114830. The content is solely the responsibility of the authors and does not necessarily represent the official views of the National Institutes of Health.

## Author contributions

FS and XJ performed electrophysiological recordings and manual neuronal reconstructions. DK supervised data analysis. JC created full-length cDNA libraries and aided in morphological recovery. LH prepared the full-length cDNA libraries for sequencing and performed sequencing and initial bioinformatics analysis under the supervision of RS. ZT and SP sustained animals’ colonies and provided experimental support. SL did the morphological data analysis. YB did the electrophysiological data analysis. DK did the transcriptomic data analysis. FS, DK, SL, YB, and EF analyzed the data and produced the figures. FS, DK, SS, CRC, PB, EF, and AST wrote the manuscript. AST, XJ, SP, and PB discussed and oversaw analysis and results. All authors revised the manuscript.

